# Bacterially produced GABA protects neurons from degeneration

**DOI:** 10.1101/711887

**Authors:** Arles Urrutia, Victor A. Garcia, Andres Fuentes, Mauricio Caneo, Marcela Legüe, Sebastián Urquiza, Juan Ugalde, Paula Burdisso, Andrea Calixto

**Affiliations:** Centro de Genómica y Bioinformática, Facultad de Ciencias, Universidad Mayor, Santiago de Chile, Chile; Centro Interdisciplinario de Neurociencias de Valparaíso, Facultad de Ciencias, Universidad de Valparaíso, Valparaíso, Chile; Instituto de Ciencias Biomédicas, Facultad de Medicina, Universidad de Chile; Instituto de Biología Molecular y Celular de Rosario (IBR-CONICET), Facultad de Ciencias Bioquímicas y Farmacéuticas, Universidad Nacional de Rosario and Plataforma Argentina de Biología Estructural y Metabolómica (PLABEM), Rosario, Santa Fe, Argentina

## Abstract

*Caenorhabditis elegans* and its cognate bacterial diet comprise a reliable, widespread model to study diet and microbiota effects on host physiology. Nonetheless, how diet influences the rate at which neurons die remains largely unknown. A number of models have been used in *C. elegans* as surrogates for neurodegeneration. One of these is a *C. elegans* strain expressing a neurotoxic allele of the MEC-4(d) DEG/ENaC channel which causes the progressive degeneration of the touch receptor neurons (TRNs). Using such model, this study evaluated the effect of various dietary bacteria on neurodegeneration dynamics. While degeneration of TRNs was steadily carried and completed at adulthood in the strain routinely used for *C. elegans* maintenance *Escherichia coli* OP50, it was significantly reduced in environmental and other laboratory bacterial strains. Strikingly, neuroprotection reached more than 40% in the *E. coli* HT115 strain. HT115 protection was long lasting well into old age of animals and not restricted to the TRNs. Small amounts of HT115 on OP50 bacteria as well as UV-killed HT115 were still sufficient to produce neuroprotection. Early growth of worms in HT115 protected neurons from degeneration during later growth in OP50. HT115 diet promoted the nuclear translocation of the DAF-16/FOXO transcription factor, a phenomenon previously reported to underlie neuroprotection caused by downregulation of the insulin receptor in this system. Moreover, a *daf-16* loss of function mutation abolishes HT115-driven neuroprotection. Comparative genomics, transcriptomics and metabolomics approaches pinpointed the neurotransmitter γ-aminobutyric acid (GABA) as a metabolite differentially produced between *E. coli* HT115 and OP50. HT115 mutant lacking glutamate decarboxylase enzyme genes (*gad*), which catalyze the conversion of GABA from glutamate, lost the ability to produce GABA and also to stop neurodegeneration. Moreover, *in situ* GABA supplementation or heterologous expression of glutamate decarboxylase in *E. coli* OP50 conferred neuroprotective activity to this strain. Specific *C. elegans* GABA transporters and receptors were required for full HT115-mediated neuroprotection. Together, these results demonstrate that bacterially produced GABA exerts an effect of neuroprotection in the host, highlighting the role of neuroactive compounds of the diet in nervous system homeostasis.

## Introduction

Intestinal microbes regulate many aspects of host physiology [1] including immune system maturation [2–4], neurodevelopment and behavior [5-7] among others. Recent reports show that in mood disorders and neurodegenerative diseases the microbiome composition and abundance is altered, and this has provided a glimpse at the role of specific bacterial metabolites with neuroactive potential in the prevention of such disorders [8-11]. However, whether bacterial metabolites directly influence neuronal degeneration and their mechanisms of action is largely unknown. The bacterivore nematode *C. elegans* continues to provide an excellent model to study the relationship between bacteria and host [12]. Both the animal and its bacterial diet are genetically tractable making them suitable for individual gene and large-scale mutation analysis. This system has been instrumental in deciphering specific metabolites from gut bacteria influencing developmental rate, fertility and ageing [13], and host factors mediating germline maintenance in response to a variety of bacterial diets [14] as well as defensive behavioral strategies against pathogens [15].

Genetically encoded prodegenerative stimuli such as a dominant mutation on *mec-4 (mec-4d)*, the gene encoding the MEC-4 degenerin have proven effective in deciphering common molecular players of neuronal degeneration in invertebrates and mammals [16-18]). The touch receptor neurons (TRNs) of *C. elegans* respond to mechanical stimuli by causing an inward Na^+^ current through the MEC-4 channel, a member of the DEG/ENaC family. Mutations near the second TM helix (A713) termed *mec-4d* cause the degeneration of the TRN and render animals insensitive to touch [19]. Necrosis of the TRNs is presumably due to Na^+^ entry as well as Ca^2+^ and reactive oxygen species (ROS) imbalance [18, 20, 21]. The degeneration of the TRN begins with the fragmentation of the axon followed by the swelling of the soma. Noteworthy, the use of this model has allowed determining interventions than can delay neurodegeneration, such as caloric restriction, antioxidant treatment and mitochondria blockage [16], as well as diapause entry [22]. In this study, we evaluated the rate of degeneration of *C. elegans* neurons in different dietary bacteria and found that specific dietary bacteria promote protection from neuronal degeneration. Combining system biology approaches coupled to genetics we discovered that γ-aminobutyric acid (GABA) produced by bacteria is protective for *C. elegans* neurons undergoing degeneration.

## Results

### Bacterial diet influences the rate at which neurons degenerate

We measured the effect of different dietary bacteria on the progression of genetically induced neuronal degeneration of the TRNs in a *C. elegans mec-4d* strain, expressing a mutant mechanosensory channel MEC-4d [19]. We previously showed that *mec-4d-*expressing Anterior Ventral Microtubule (AVM) touch neuron dies in a stereotyped fashion and defined window of time when animals feed on the standard laboratory *E. coli* OP50 diet [16]. Right after hatching, *mec-4d* mutant animals were fed different bacteria and the AVM neuronal integrity was quantified in adulthood, 72 hours later. The dietary bacteria used were *E. coli* OP50, *E. coli* HT115, *E. coli* K12, *Comamonas aquatica, Comamonas testosteroni, Bacillus megaterium* and the mild pathogenic *P. aeruginosa* PAO1. In accordance to our previous reports, neurodegeneration steadily occurred when feeding in *E. coli* OP50 on which only a very low percentage of worms (1-3%) maintained AVM axons after three days (**Figure 1A)**. Notably, although neurodegeneration occurred in *E. coli* B and *C. testosteroni* similarly to when feeding on *E. coli* OP50, the bacteria *E. coli* HT115, *E. coli* K12, *C. aquatica, B. megaterium* and *P. aeruginosa* gave significant protection (**Figure 1B**). *E. coli* HT115 was the most protective, with over 40% of wild type axons 72 hours after hatching, compared to less than 2% in *E. coli* OP50 (**Figure 1B, D**). The broad difference on neuronal integrity in *mec-4d* worm populations feeding on *E. coli* OP50 or HT115 can be observed in **Figure 1C**.

**Figure 1.**
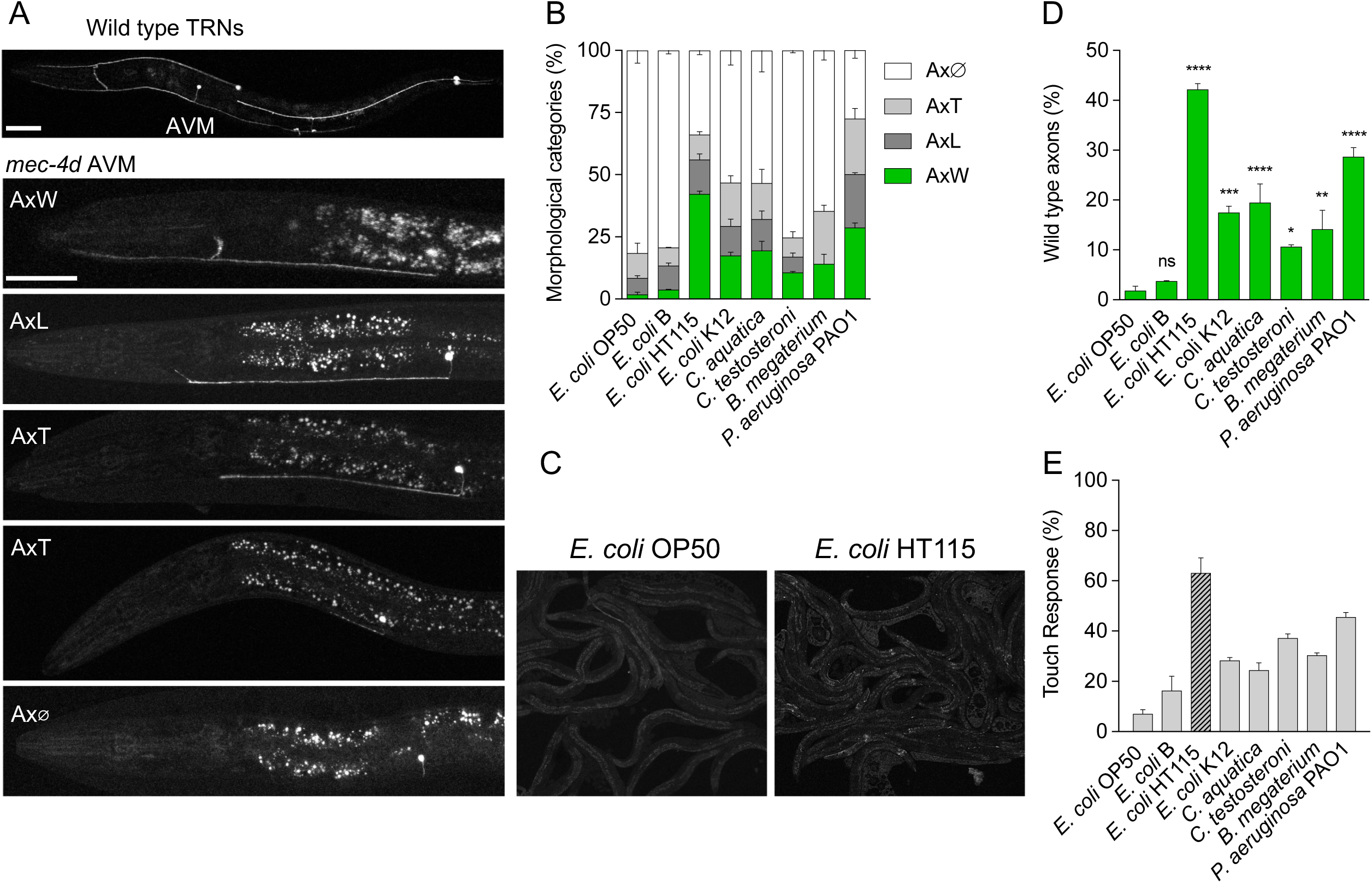
Dietary bacteria determine the rate of neuronal degeneration. **A**. Touch receptor neurons expressing GFP in wild type and *mec-4d* worms. The latter shows the stereotypical progression of AVM degeneration of *mec-4d* mutants and constitutes the axonal categories assessed during the experiment shown in B. Scale bars represent 20 μm. **B**. Percentage of all morphological axonal categories in *mec-4d* worms after 72 hours of growth in different bacterial strains. **C**. Fluorescence microscopy fields of GFP-expressing *mec-4d* worms raised in the indicated *E. coli* strains, comparing the presence of AVM axons on each preparation at 10X magnification. **D.** Percentage of wild type axons in the experiment shown in B. **E**. Percentage of touch responsiveness of animals after growth in the different dietary bacteria.

TRNs are neurons expressing receptors of gentle mechanical stimuli [23]. Hence, we determined the response to gentle touch in worms fed the different strains to test whether morphological protection shown in **Figure 1B** translate into functional responses. **Figure 1E** shows that the number of responses in worms correlates with the morphological categories AxW and AxL, the two axonal categories defined as functional in previous work [22].

### Bacterial components promote neuroprotection

Phenotypical outcomes mediated by intestinal bacteria can be a result of either a modulation of host physiology by interspecies live interactions (i.e. bacterial colonization) or by the exposure of the host to a bacterial metabolite. The first one requires bacteria to be alive in the intestine, while the second does not. To distinguish between these two possibilities, we fed *mec-4d* animals with UV-killed HT115 bacteria; the most protective among those tested and scored the AVM integrity at 72 hours. Dead bacteria protected to the same extent as live bacteria (**Figure 2A**). This indicates that protective molecules of *E. coli* HT115 bacteria are produced in bacteria prior to the exposure to the animals and thus, neuronal protection is independent of the induction of bacterial responses by the interaction between the HT115 bacteria and the host. Furthermore, worms raised in dead HT115 bacteria cultivated to different optical densities (OD) displayed the same levels of neuroprotection, suggesting that the protective factors are present in the bacteria during all phases of the growth curve (**S1A-B Figures**).

**Figure 2.**
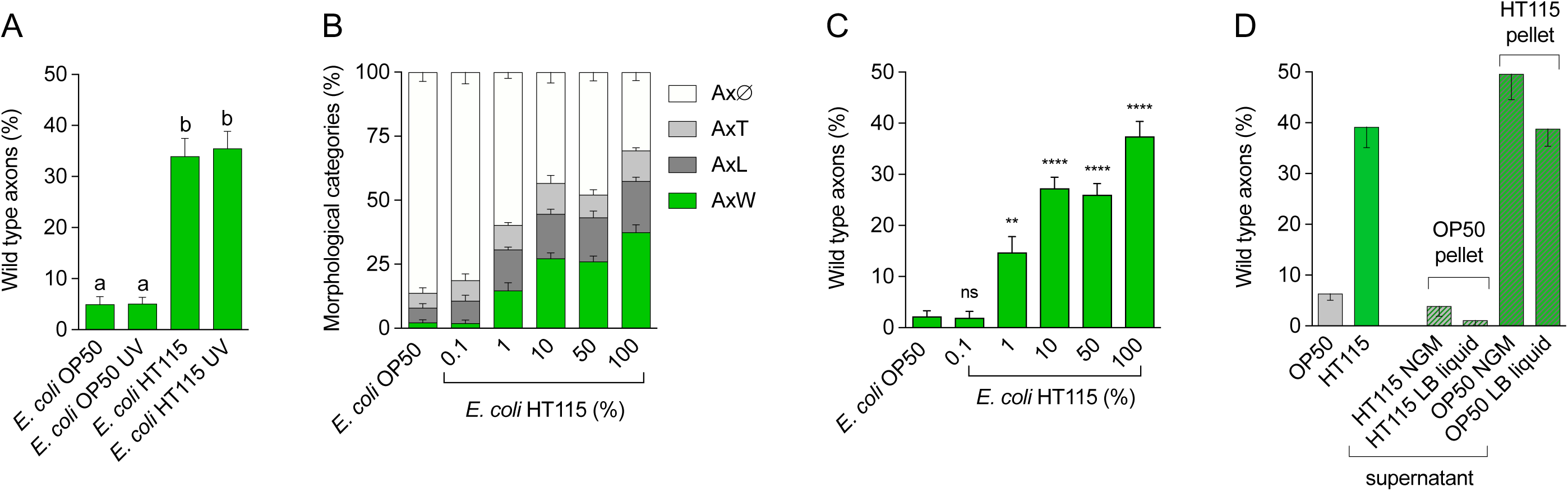
Bacterial components have neuroprotective activity. **A.** Axonal categories in worms raised in the indicated live or UV-killed bacterial strains. **B-C**. All axonal categories (**B**) or wild type (**C**) axons in worms rose in different proportions of UV-killed *E. coli* HT115 on UV-killed OP50. **D.** Wild type axons of *mec-4d* animals fed with *E. coli* HT115 or OP50 bacterial pellet supplemented with supernatant of OP50 or HT115. **E-F**. All axonal categories (**E**) and wild type axons (**F**) of *mec-4d* animals fed with *E. coli* OP50 supplemented with reconstituted extracts from *E. coli* HT115.

The large difference in neuroprotection between *E. coli* OP50 and HT115 strains raises two possibilities; one being that *E. coli* OP50 actively promotes the degeneration of the neuron and the other that HT115 has a protective effect. To discern this matter, we fed worms with a mix of U.V killed *E. coli* HT115 and OP50 in different proportions (**Figures 2B-C**). A 1/100 (1%) dilution of *E. coli* HT115 was sufficient to protect AVM neurons significantly more than undiluted OP50. This strongly suggested that *E. coli* HT115 produces a neuroprotective compound needed in small amounts.

To test whether the neuroprotective molecules are being secreted by the bacteria, we separated the supernatant of both bacterial strains from their pellets by centrifugation and mixed the supernatant of *E. coli* HT115 with OP50 pellet and vice versa (see Materials and Methods for details). *E. coli* HT115 supernatant was not capable of providing protective activity when mixed with *E. coli* OP50 pellet (**Figure 2D**). This suggests that the protective factor is not secreted, or that the amounts contained in the supernatant are not sufficient for protection. As expected, *E. coli* OP50 supernatant did not altered the protection pattern of *E. coli* HT115.

### *E. coli* HT115 diet promotes long-term protection of mechanoreceptors and interneurons of the touch receptor circuit

*E. coli* HT115 showed to be neuroprotective throughout the development of the animal and into young adulthood (**Figure 1B**). We explored if AVM neurons are still protected after maturity of worms. To that end we fed newly hatched *mec-4d* animals with *E. coli* HT115 and scored their neuronal integrity every 24 hours for 168 hours. While on *E. coli* OP50 all animals had degenerated neurons at the final time point, on HT115 food 25% of animals had wild type AVM axons (**Figures 3A-C**), confirming the ability of HT115 to significantly protect at later life stages. Noteworthy, between 12 and 24 hours after hatching in HT115 there was a statistically significant rise in AxW axons (**Figure 3C**), suggesting that neurons could be growing after an initial truncation. To assess this, we followed individual animals in a longitudinal fashion on *E. coli* HT115 and scored the neuronal integrity of each nematode every 24 hours for 3 days. We scored axons separately according to their initial and final morphology and classified axonal outcome as *Protection* when the morphology of axons did not change from truncated or wild type, *Degeneration* when axonal morphology changed from AxT to Ax□ or was maintained as Ax□. Finally, *Regeneration* refers to axon growth from truncations to wild type. While the most prevalent category is Protection (40%), 30% of axons regenerated between 24 to 72 hours after hatching on *E. coli* HT115 (**Figure 3D)**. This suggests that under HT115 protective conditions a portion of neurons can repair broken axons.

**Figure 3.**
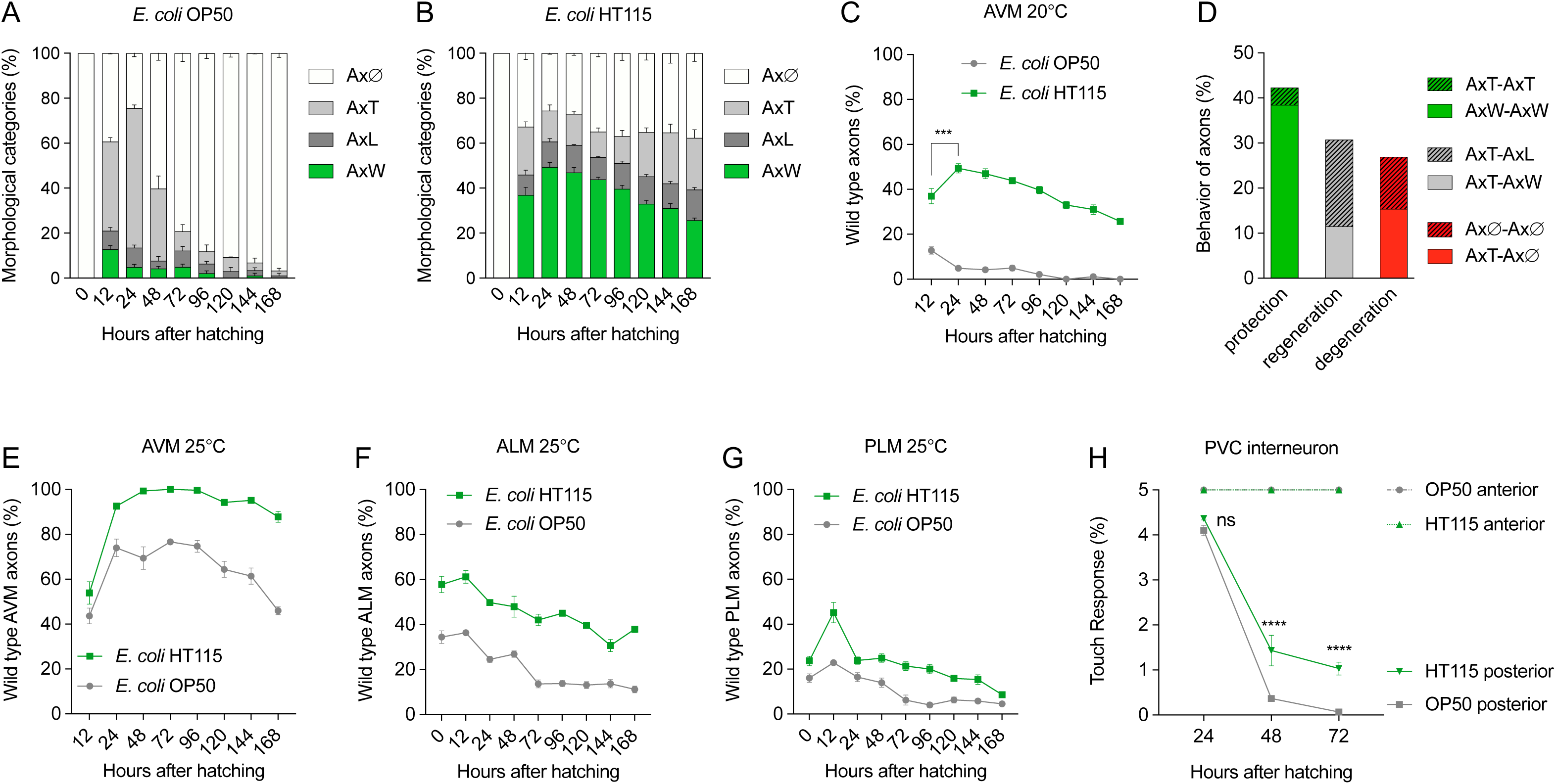
Neuroprotection induced by dietary *E. coli* HT115 is long lasting and extended to other neuronal types. Time course of axonal categories of worms fed with *E. coli* OP50 (**A**) or HT115 (**B**) for 168 hours. **C**. Percentage of wild-type axons in **A** and **B. D.** Proportion of axonal categories in longitudinal assays. *Protection* (green) indicates axons that were not degenerated over time regardless of the initial category, with the exception of AxØ. *Regeneration* (gray) accounts for axons that grew in size over time. *Degeneration* (red) accounts for axons that degenerated over time. Determination of wild type axons in AVM (**E**), ALM (**F**), PLM (**G**) and PVC (**H**) neurons of worms fed *E. coli* OP50 or HT115.

Next, we explored whether other neurons of the touch circuit are protected from degeneration in *E. coli* HT115 diet. It has been already reported that at hatching, four embryonic TRN, two Anterior Lateral Microtubule (ALMs) and two Posterior Lateral Microtubule (PLM) neurons have already degenerated when growing at 20°C in this model [16, 24]. At 25°C however, degeneration proceeds at a slower rate [16, 22]. To analyze the degeneration rate of ALM and PLM neurons, L4 animals were grown at 25°C and their progenies synchronized at birth. The neuronal integrity of ALM, PLM and AVM cells was assessed at 12, 24 and every 24 hours after birth until 168 hours at 25°C. The percentage of wild type neurons in *E. coli* HT115 diet is significantly higher than that in OP50 diet throughout the temporal course for all three neurons (**Figures 3E-G**, full morphological characterization is shown in **S2A-F Figures)**.

We tested next whether *E. coli* HT115 was capable of protecting the forward command interneuron PVC expressing the *deg-1* prodegenerative stimulus. *deg-1(u38)* animals progressively lose the ability to respond to posterior touch due to the time dependent degeneration of the PVC interneuron [25]. We tested the posterior touch response of *deg-1* animals during development feeding on *E. coli* OP50 and HT115. *E. coli* HT115 promotes a larger functional response than *E. coli* OP50 suggesting that this neuron is also protected (**Figure 3H**). Taken together, these results demonstrate that HT115 diet is protective over different neuronal types undergoing degeneration.

### Early exposure of animals to *E. coli* HT115 is sufficient for neuronal protection

*Ad libitum* feeding on *E. coli* HT115 protected *mec-4d* expressing neurons from degeneration for long periods of time. We sought to investigate if a constant stimulus provided by the HT115 metabolite is required to achieve neuroprotection or if an early, discrete time-lapse exposure to the diet is sufficient. We fed animals for the first 6 hours after hatching (previous to the birth of the AVM) and for 12 hours after hatching (at birth of the neuron) with *E. coli* HT115 and immediately switched to *E. coli* OP50. We scored the neuronal morphology 12, 24, 48 and 72 hours post hatching (**S3A, D Figures**). In parallel, both diets were fed *ad libitum* as controls. Strikingly, animals that ingested *E. coli* HT115 for only 6 hours showed a significantly larger number of wild type neurons at 72 hours (14.3 %) than animals continuously fed OP50 (3.6 %, **Figure 4A**), and had more axons in the other categories (**S3A, C Figures**). Feeding HT115 to *mec-4d* animals for the first 12 hours after hatching conferred a significantly larger protection than feeding HT115 for only 6 hours (**Figure 4A**). These results show that although early short exposures are not equally protective than a permanent HT115 diet, they do have a long-lasting effect in neurons compared to an uninterrupted diet of *E. coli* OP50.

**Figure 4.**
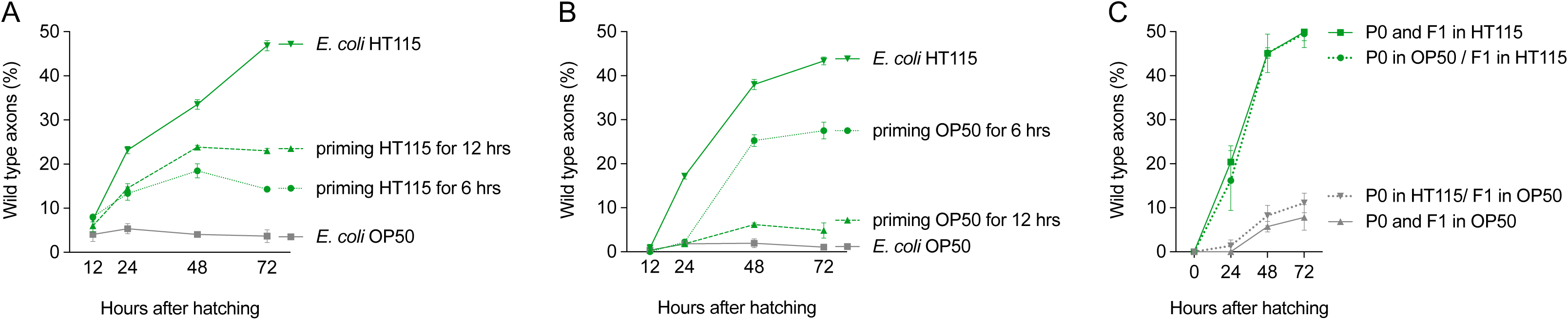
Diet of *E. coli* HT115 at early stages of life is necessary and sufficient to confer neuroprotection. **A-B** Percent of wild type axons of animals fed for 6 and 12 hours with (**A**) *E. coli* HT115 or (**B**) *E. coli* OP50 and then changed to OP50 or HT115 diet respectively. (**C**) Percent of wild type axons of animals feeding on either *E. coli* OP50 or HT115 whose parents were fed on either diet.

We then tested the effect of early exposure to non-protective bacteria. We fed *mec-4d* animals for 6 and 12 hours with *E. coli* OP50 and then changed them to HT115. The morphology of AVM neurons was scored at 12, 24, 48 and 72 hours post hatching. 6 hours of *E. coli* OP50 exposure did not prevent HT115 from protecting AVM neurons later in adulthood. Exposure for 12 hours however precludes protection of the AVM (**Figure 4B**). This suggests that the time between the first 6 and 12 hours of development is crucial for the protective effect to take place.

In *C. elegans*, some dietary bacteria-induced traits show heritable properties [15]. Therefore, we tested whether neuronal protection could be inherited. Animals were fed either *E. coli* OP50 or HT115 from birth and their F1 progeny passed to OP50 or HT115. Neuronal integrity of descendants was tested in a time course fashion. One generation of parental feeding on HT115 did not improve neuronal protection in the progeny feeding on OP50, nor did *E. coli* food preclude protection of F1 feeding on HT115 (**Figure 4C**). This result indicates that the protective effect of *E. coli* HT115 is not transmitted intergenerationally.

### *E. coli* HT115 protection requires DAF-16 signaling

We explored the role of the Insulin/IGF-1-like signaling (IIS) pathway, a well-described and conserved signaling cascade acting systemically. In *C. elegans*, downregulation of the insulin receptor DAF-2 promotes neuroprotection [16]. We investigated whether *E. coli* HT115 neuroprotective effect also involved the IIS. First, we fed *daf-2ts; mec-4d* animals with *E. coli* HT115 at 25°C. This strain expresses a DAF-2 protein version that is unstable at 25°C. Next, we scored the neuronal integrity of ALM, PLM and AVM neurons in a time dependent fashion. ALM and PLM protection in *E. coli* HT115 was not further increased by the elimination of *daf-2* (**Figures 5A-B**, neuronal integrity on *E. coli* OP50 of ALM, PLM and AVM at 25°C in **S4A-C Figures**), suggesting that HT115-mediated protection involves the downregulation of the DAF-2 pathway. Owing to the cumulative protection effects of temperature and diet, *mec-4d* AVM neurons at 25°C on HT115 diet reached almost 100% of wild type axons and the *daf-2* mutation maintained maximum protection (**Figure 5C)**.

**Figure 5.**
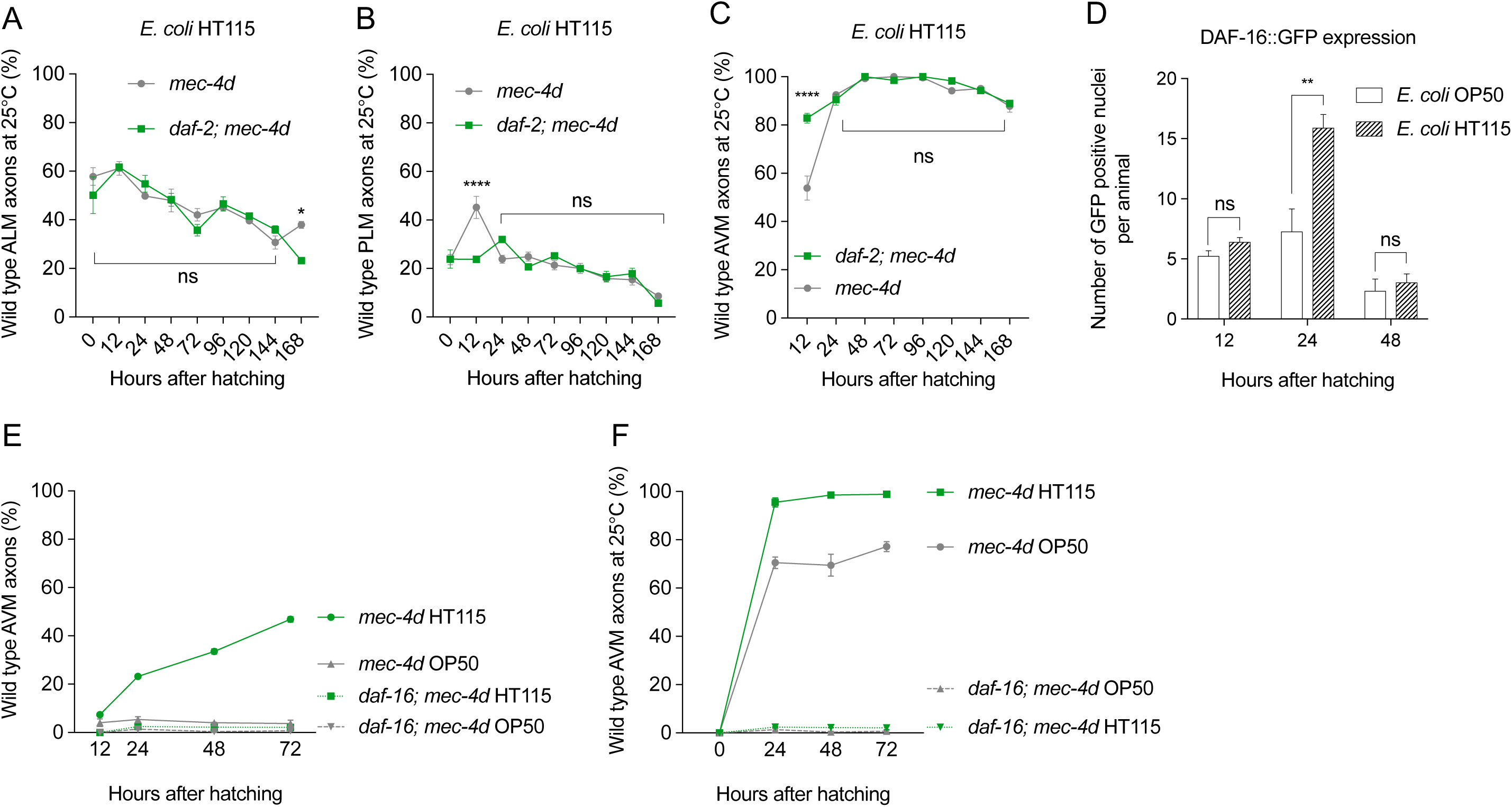
Effect of downregulation of the insulin pathway on *E. coli* HT115 mediated neuroprotection. **A-C** Neuronal integrity of ALM (**A**), PLM (**B**) and AVM (**C**) neurons of *daf-2(ts*); *mec-4d* animals feeding on HT115 food. **D**. Number of GFP positive nuclei in DAF-16::GFP animals feeding on *E. coli* OP50 or HT115 during development. **E-F**. Time course of neuronal degeneration in *daf-16; mec-4d* animals at 20°C (**E**) and 25°C (**F**).

Because DAF-2 downregulation causes the translocation of DAF-16 to the nuclei [26], we tested whether HT115 diet promoted nuclear translocation of GFP to nuclei in a DAF-16::GFP expressing strain (CF1139 strain, see Materials and Methods). We fed CF1139 animals with *E. coli* OP50 and HT115 and compared the number of fluorescent nuclei at 12, 24 and 48 hours after hatching on each diet. *E. coli* HT115 promoted a significantly higher translocation of DAF-16 compared to *E. coli* OP50 at 24 hours only, returning to basal levels at 48 hours (**Figure 5D**).

Next, we directly assessed the involvement of DAF-16 in the neuroprotection effect of *E. coli* HT115 by scoring neuronal integrity of the TRNs in *daf-16; mec-4d* animals feeding with HT115 at 20°C and 25°C. Both in *E. coli* OP50 and HT115 most TRNs types were absent at birth in either temperature, with a marginal presence of AVMs, which also rapidly underwent degeneration in the *daf-16* mutant (**Figure 5E-F**, all categories in *E. coli* OP50 and HT115 at 20°C and 25°C in **S4D-G Figures)**. This demonstrated that DAF-16 is required for the neuroprotection effect of the HT115 diet. Noteworthy, the *daf-16* mutation completely abolished the protection of TRNs previously observed at 25°C in *mec-4d* background [22] in both bacteria, suggesting that the *daf-16* mutation significantly lowers the threshold for neurodegenerative stimuli. The effect of increased degeneration in *daf-16; mec-4d* though much more dramatic in *E. coli* HT115 is also observed in OP50. We assessed the effect of *daf-16* mutations in normal development and integrity of the TRNs in the strain WCH40 [*daf-16*(*m27*); *uIs31*(*P*_*mec-17*_*mec-17::gfp*)] expressing GFP in all the TRNs. DAF-16 loss alone did not cause an observable effect in the morphology of the TRNs (**S4H Figure**). Taken together, these results show that DAF-16 is necessary for the protective dietary protection mediated by *E. coli* HT115.

### Identification of uniquely expressed genes on neuroprotective bacteria

To identify the bacterial molecule(s) conferring neuroprotection we looked for differences in the genomes and transcriptomes of the two *E. coli* strains. We reasoned that genes important for neuronal protection would be uniquely expressed or upregulated in *E. coli* HT115 compared to OP50. We first sequenced the genomes of *E. coli* HT115 and OP50 using the Illumina MiSeq platform. Unique and shared genes from each strain are represented in **Figure 6A** and unique genes are detailed in **Supplementary File 1**. Gene expression of each bacterium as well as differential expression determined using DeSeq2 are reported in **Supplementary File 2**. We grouped unique and common genes according to their expression levels (**Supplementary Table 1).** Expression levels of the top highly expressed genes in both strains are shown in **Figure 6B**. Only two genes unique to *E. coli* HT115 are highly expressed: *gadA* (4368 TPM) encoding the glutamic acid decarboxylase and *cspB* (2655 TPM) encoding a cold shock-like protein.

**Table 1.**
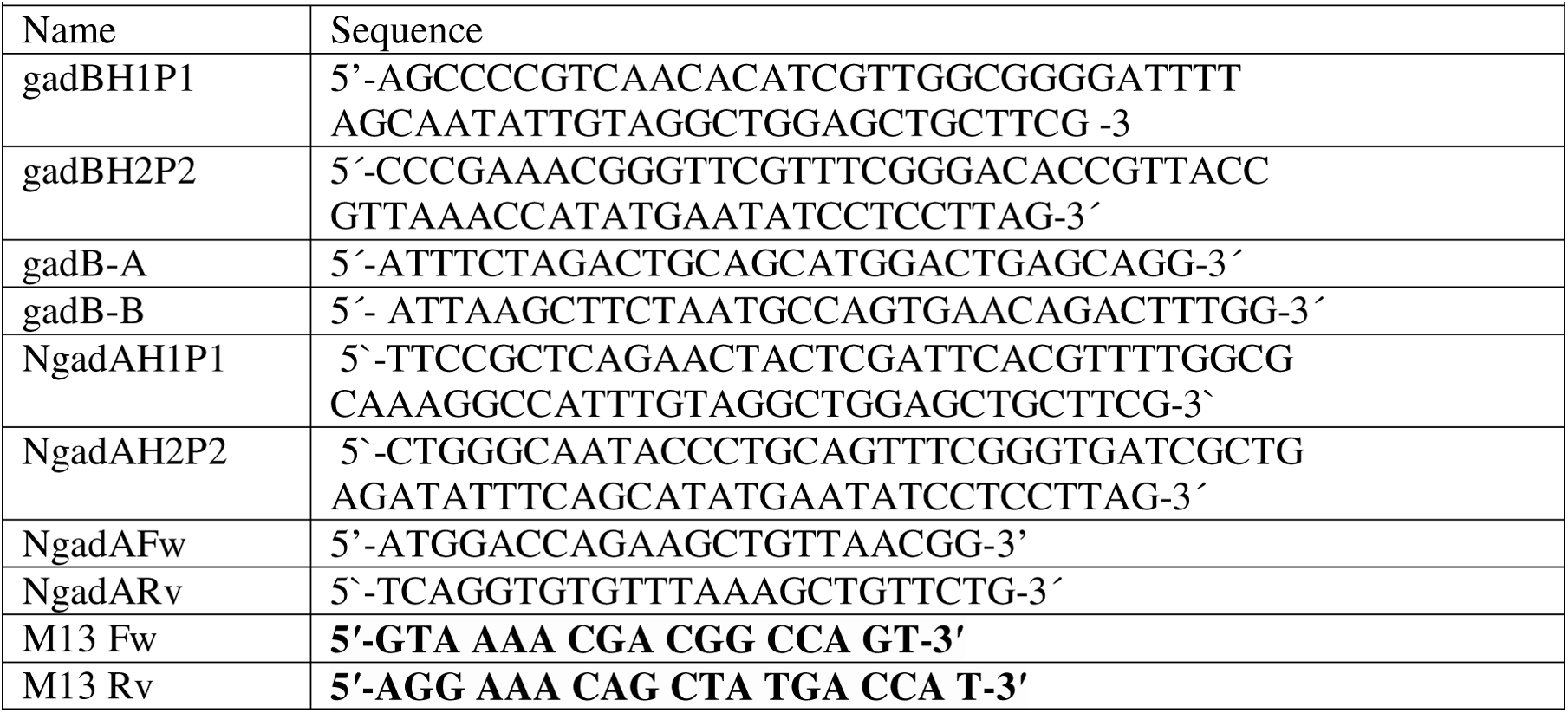
Primers used for the construction of the *E. coli* HT115 Δ*gadB*::*cat*/Δ*gadA*::*kan* double mutant strain.

**Figure 6.**
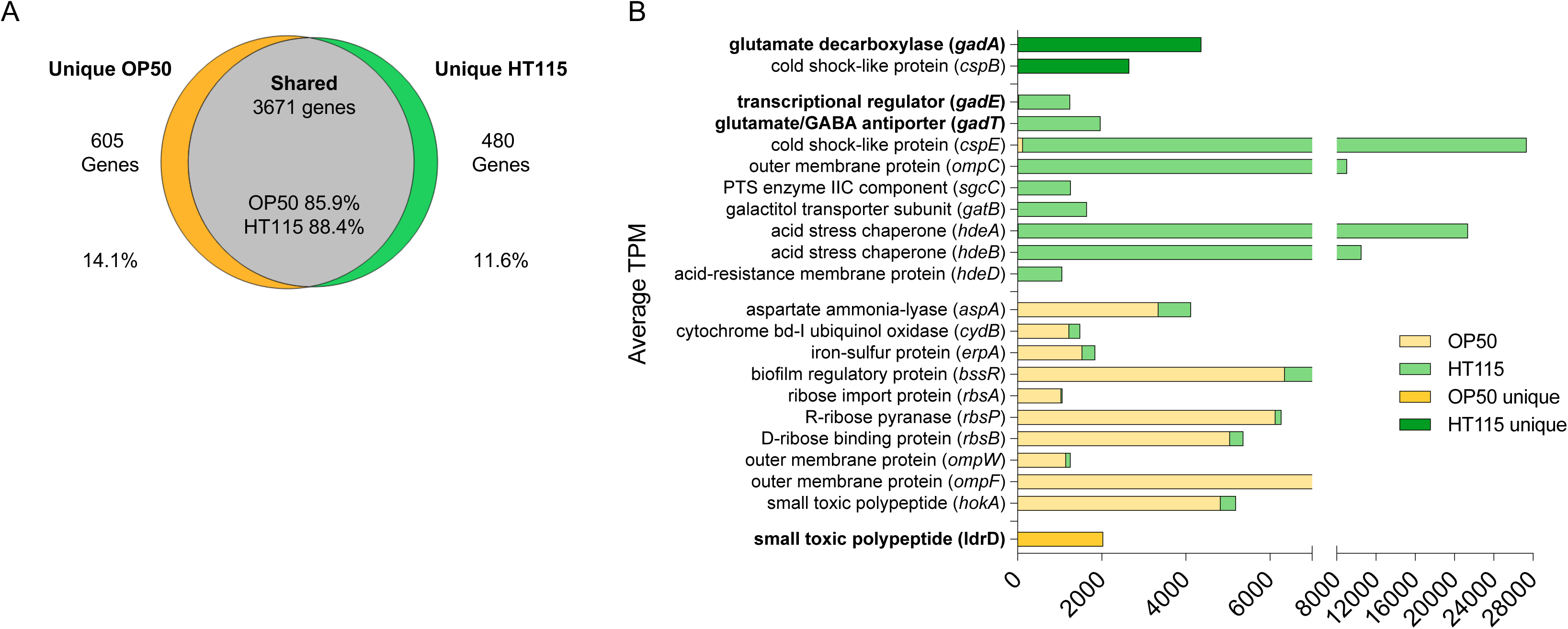
Glutamate decarboxylase enzyme is uniquely expressed in *E. coli* HT115. **A**. Percent of unique and shared genes between the two bacterial genomes. **B**. Average Transcripts per million (TPM) of highly and uniquely expressed genes in *E. coli* HT115 or OP50 (bold) and the 10 genes with the largest fold change of expression in both bacteria compared to each other.

Interestingly, *gadA* that encodes glutamate decarboxylase (GAD) the enzyme catalyzing the conversion of glutamic acid to γ-amino butyric acid (GABA) is the *E. coli* HT115 gene with the highest expression among unique genes. For this reason, we chose GAD as our first candidate enzyme to evaluate neuroprotective metabolite production. We also looked at other enzymes involved in the utilization or export of GABA that were overexpressed in *E. coli* HT115. The glutamate/GABA antiporter *gadC* is among the shared genes highly expressed in HT115 versus OP50 (**Figure 6B**) with a log2 fold change (L_2_FC) of 9.8 over OP50, suggesting that GABA can be exported to the periplasma of bacteria. The enzyme that metabolizes GABA, γ-aminobutyric transaminase (*gabT*) is expressed at medium levels (18 TPM in average) and a L_2_FC of 2.5; succinic semialdehyde dehydrogenase (*gabD*) is expressed at low levels (8.9 TPM in average) and a L_2_FC of 2.5. The transcriptional regulators genes *rcsB* and *gadE*, required for the activation of the *gad* operon, are among the genes expressed at medium (315 TPM) and high (1219 TPM) levels in HT115, respectively. The latest has a L_2_FC of 5.2 over OP50 (**Supplementary File 2**). This suggests that enzymes and metabolites involved in the pathway of GABA production and utilization are good candidate neuroprotective players.

### GAD and GABA are required for *E. coli* HT115 neuroprotection and GAD activity correlates with neuroprotection by other bacteria

To test the role of GAD and its product GABA in neuroprotection, we first generated a *gad* null mutant of *E. coli* HT115 by homologous recombination (HT115Δ*gad*, details in Materials and Methods). To demonstrate that HT115Δ*gad* lacked GAD activity, we used a colorimetric assay based on pH elevation given by the conversion of glutamate to GABA [27, 28]. As expected, wild type *E. coli* HT115 raised the pH of the solution while neither HT115Δ*gad* nor OP50 were able to do so. To confirm that a raise in pH is due to the expression of GAD, we transformed *E. coli* OP50 with a plasmid expressing *gadA* (pG*gadA*). *E. coli* OP50 pG*gadA* supplemented with glutamate showed potent enzymatic activity, rising the pH of solution above HT115 levels (**Figure 7A**). Additionally, we directly measured GABA production in the three bacteria using the GABase test [29]. *E. coli* HT115 pellet had the highest GABA levels while OP50 and HT115Δ*gad* were indistinguishable from each other (**Figure 7B**). This demonstrates that GABA is being produced in *E. coli* HT115 and not in OP50 or the HT115 Δ*gad* strain.

**Figure 7.**
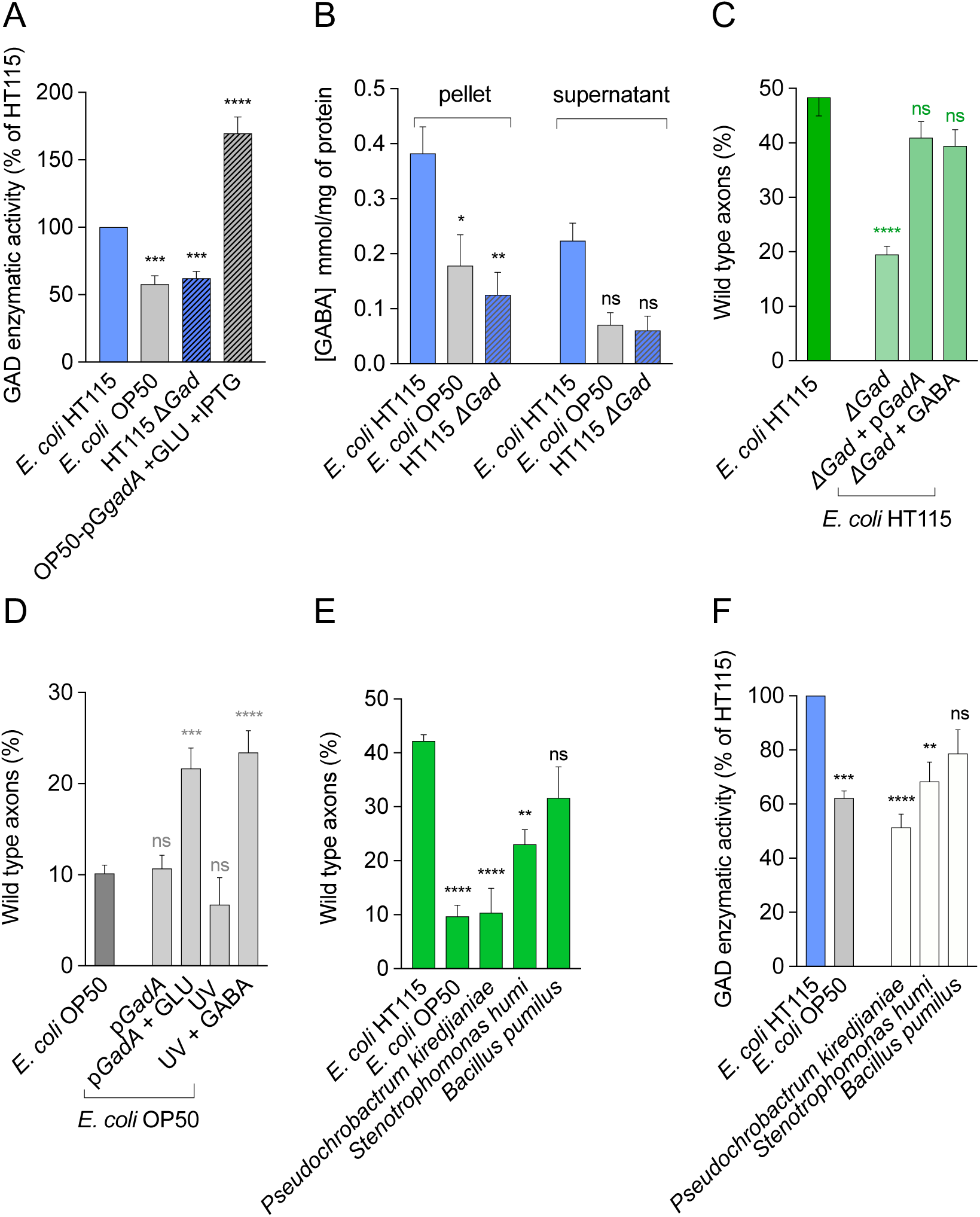
Bacterial GABA is crucial for neuroprotection. **A**. Measurements of GAD enzyme activity normalized as a percentage of HT115 GAD activity in wild type, mutant, and transformed bacterial strains. **B**. GABA concentration in the pellet and supernatant of HT115 wild type and Δ*gad* mutant. **C-D**. Percent of wild type axons in wild type and Δ*gad* mutant HT115 strain (**C**) and wild type OP50 supplemented with GABA and genetically transformed with the pG*gadA* plasmid (**D**). **E-F**. Neuroprotection and GAD activity of bacterial isolates from a natural microbiota. **E**. Percent of wild type axons of animals fed with wild bacterial isolates. **F**. Measurements of GAD enzyme activity normalized as a percentage of *E. coli* HT115.

Secondly, we fed *mec-4d* animals with *E. coli* HT115Δ*gad* and scored its protective potential at 72 hours. HT115Δ*gad* was not able to protect degenerating AVM neurons, showing a significantly reduction of wild type axons compared to wild type strains (**Figure 7C**). This shows that GAD activity plays a pivotal role in the protection conferred by HT115 bacteria. Moreover, plasmid pG*gadA* was able to rescue protective potential in null mutant HT115Δ*gad*. Additionally, dietary supplementation of HT115Δ*gad* with 2 mM of GABA was sufficient to provide neuroprotection (**Figure 7C)**. Finally, we fed *mec-4d* with *E. coli* OP50 pG*gadA* to test whether *gadA* is sufficient to provide protective activity in the presence or absence of glutamic acid, the substrate of GAD. While *E. coli* OP50 pG*gadA* alone was not sufficient to increase wild type axons incidence, glutamate addition to the bacterial culture protection was significantly increased compared to *E. coli* OP50 pG*gadA* and *E. coli* OP50 wild type (Figure 7D). This is coherent with increased GAD activity of *E. coli* OP50 supplemented with pG*gadA* shown in **Figure 7A**. Furthermore, supplementation of HT115Δ*gad* with 2 mM of GABA was sufficient to provide neuroprotection (**Figure 7C)**. Importantly, *E. coli* OP50 supplemented with GABA also protected *mec-4d* neurons (**Figure 7D**). Taken together these results show that GAD and its product GABA play an important role in *E. coli* HT115 mediated neuroprotection.

Because other bacteria also protected degenerating neurons (**Figure 1B**), we evaluated whether GAD activity was detectable in these strains. We measured GAD activity normalized against *E. coli* HT115 and showed that GAD expression is correlated with neuroprotective activity in all strains (**S5 Figure**). This suggests that GAD activity also plays a role in the neuroprotective capacity of other bacteria.

As a soil nematode, *C. elegans* feeds on a large range of bacteria in its natural environment [30]. To test whether soil bacteria found in wild *C. elegans* were capable of providing neuronal protection, we selected three bacterial species previously co-isolated with wild *C. elegans* from soil by our group, namely *Pseudochrobactrum kiredjianiae, Stenotrophomonas humi* and *Bacillus pumilus*. We fed *C. elegans* with each bacterium and scored their neuronal integrity after 72 hours comparing it to that in *E. coli* HT115 and OP50. *P. kiredjianiae* and *S. humi* did not promote neuroprotection while *B. pumilus* protected to a similar extent than HT115 (**Figure 7E**, all categories in **S7 Figure**). GAD activity of the isolates was measured and compared to that provided by *E. coli* OP50 and HT115 (**Figure 7F**). Consistent with the degree of neuroprotection produced by these bacteria, *P. kiredjianiae* and *S. humi* did not show significant GAD activity while in *B. pumilus* the GAD activity was similar to HT115. These results show that GAD activity is correlated with neuroprotection in different bacteria and supports the previous evidence that bacterial GAD enzyme and its product GABA are key for neuroprotection.

### Identification of metabolites expressed in neuroprotective conditions

To unbiasedly identify potentially neuroprotective metabolites produced by the strain HT115 but absent in the non-protective HT115Δ*gad* and *E. coli* OP50 strains, we implemented a non-targeted metabonomics approach using 1H Nuclear Magnetic Resonance. A total of 24 extract samples were prepared and analyzed (8 of each strain). To evaluate the global metabolic profile of the three bacterial strains, we performed a principal component analysis (PCA) of binned 1H NMR datasets. As we expected, all three bacteria strains were metabolically different with *E. coli* HT115 and HT115 Δ*gad* closer than *E. coli* OP50 in the metabolic space **(S6A Figure)**. Metabolites related with neuroprotection were evaluated by Orthogonal Projections to Latent Structures Discriminant Analysis (OPLS-DA), first comparing *E. coli* HT115 strain against HT115 Δ*gad* (**Figure 8A**), and secondly *E. coli* HT115 against HT115 Δ*gad* and OP50 (**Figure 8D**). OPLS-DA models were validated by 200 permutations (**S6B-C Figure**). Discriminant analysis in HT115 vs HT115 Δ*gad* revealed intergroup metabolic differences. The discriminant upregulated metabolites in wild type HT115 were GABA, lactate, sucrose and maltose, while in HT115 Δ*gad* were glutamate and putrescine (**Figure 8B-C, Supplementary Table 2, S7 Figure**). This is coherent with the accumulation of the GAD substrate, glutamate, given the absence of the enzyme on HT115 Δ*gad*. Notably, the comparison between *E. coli* HT115 strain against HT115 Δ*gad* and *E. coli* OP50 revealed intergroup metabolic differences GABA, lactate, sucrose and maltose were highly expressed in the neuroprotective strain, while there were no discriminant metabolites found in higher levels in OP50 and HT115 Δ*gad* (**Figure 8E-F, Supplementary Table 2**). Overall, in strong agreement with our genetic and chemical complementation approach, these results further indicated that GABA is the metabolite playing a central role in HT115-conferred neuroprotection.

**Figure 8.**
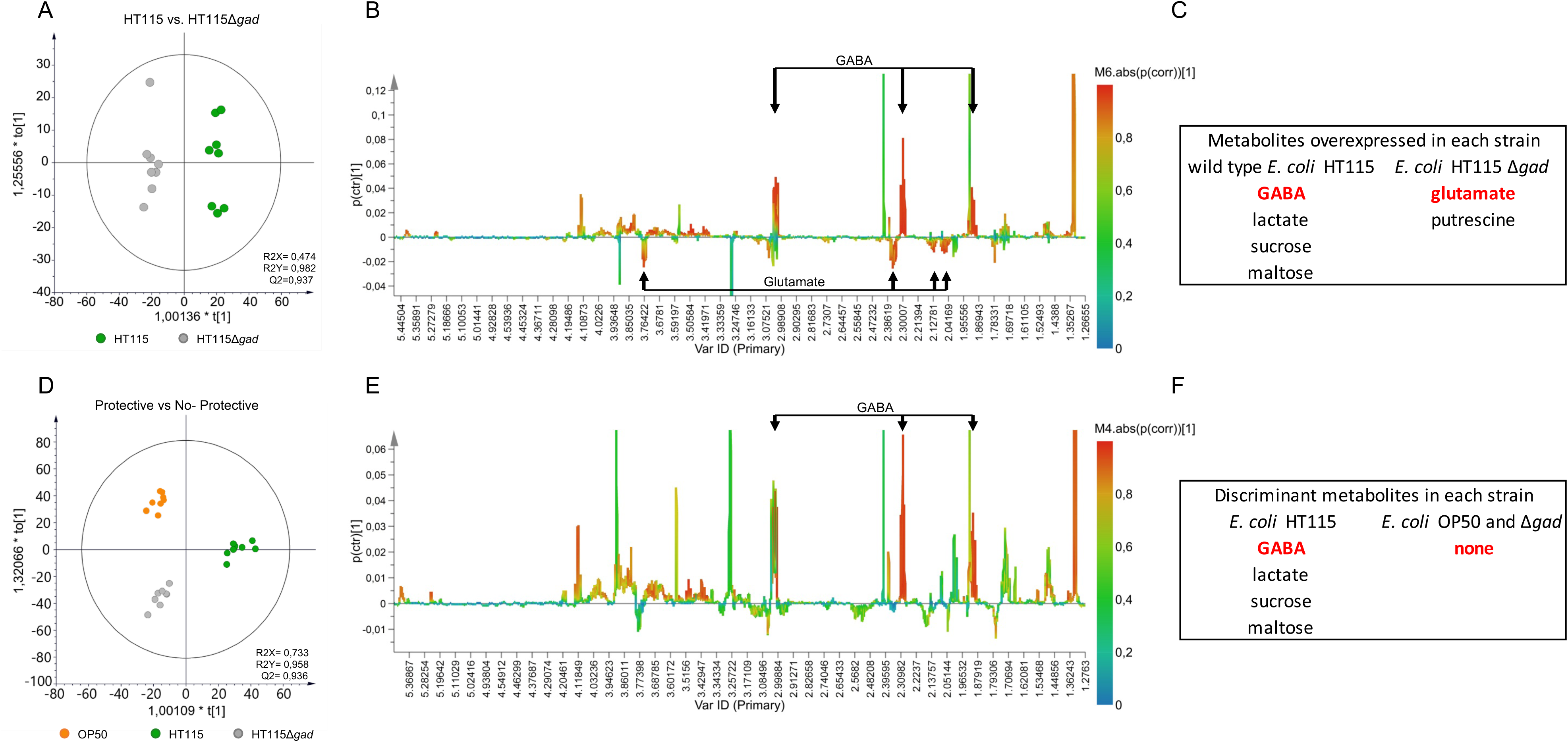
Metabolomics analysis of neuroprotective and non-protective bacteria. **A-D.** Orthogonal Projections to Latent Structures Discriminant Analysis (OPLS-DA) score plot of protective *E. coli* HT115 wild type (blue) and non-protective and *E. coli* HT115Δ*gad* (light gray) (**A**) and *E. coli* HT115 wild type (blue) and non-protective, *E. coli* HT115Δ*gad* (light gray) and OP50 (dark gray) (**D**). **B-E.** OPLS-DA S-line plots with pairwise comparison of data from NMR spectra obtained comparing *E. coli* HT115 strain against HT115 Δ*gad* **(B)** and E. coli HT115 strain against HT115 Δ*gad* and *E. coli* OP50 **(E)**. Colors are associated with the correlation of metabolites characterizing the 1H NMR data for the class of interest with the scale shown on the right-hand side of the plot. In B GABA and glutamate signals are shown. **C-F**. Tables indicate which metabolites are differentially expressed in each strain.

### Host GABA receptors and transporters are required for full HT115 bacteria mediated neuroprotection

To discern whether systemic or neuronal GABA transport was implicated in neuroprotection, we silenced the expression of a number of candidate solute transporters (*unc-47, snf-5*) and GABA receptors (*gab-1, lgc-37, unc-49*) using RNA interference. To distinguish a systemic from a touch cell specific requirement for these effectors, we used *mec-4d* animals (where neuronal RNAi is inefficient) and *mec-4d* animals sensitive to RNAi only in the TRN (WCH6, [16]. We fed dsRNA of the selected effectors to both strains and the neuronal morphology was assessed at 72 hours. *lgc-37, snf-5, unc-47* and *gab-1* dsRNA expressing bacteria caused a discrete but significant decrease in neuronal protection in the systemic RNAi strain but not in the TRN specific strain (**Figures 9A-B**), suggesting these genes act in non-neuronal tissues to mediate neuroprotection.

**Figure 9.**
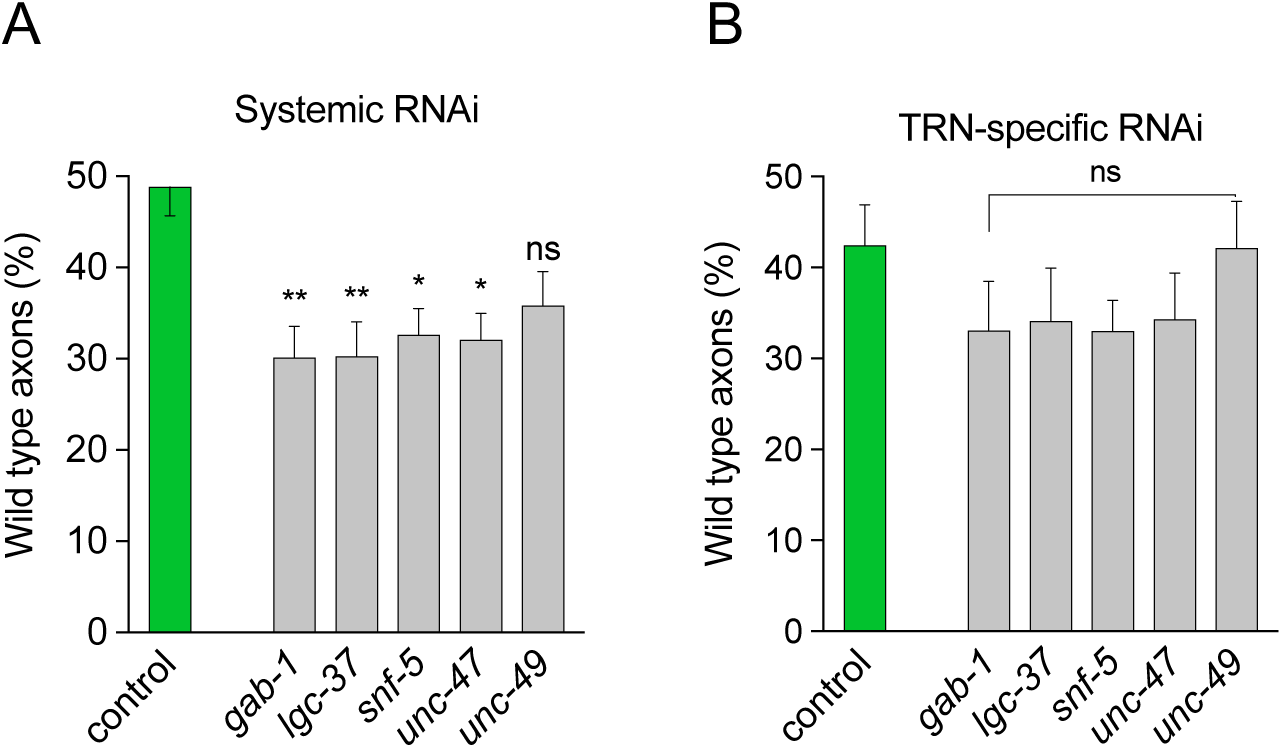
Effect of silencing GABA effectors in *C. elegans* on neuroprotection. Morphological integrity of AVM in animals feeding on dsRNA expressing bacteria for the indicated genes in a systemic (A) and TRN autonomous (B) RNAi strain.

## Discussion

The relationship between bacteria and host affects virtually every studied aspect of an animal’s physiology. However, whether bacteria and their metabolites can modulate neuronal degeneration is not known. In this work we show that bacterial diet dramatically influences neuronal outcomes in a *C. elegans* model of neurotoxic death triggered by the MEC-4d degenerin. *E. coli* HT115, a derivative of the K12 strain, is the most protective of all bacteria tested, which included non-pathogenic laboratory bacteria, mild pathogens and natural commensal bacteria. We found that this bacterium protects embryonic and postembryonic TRNs as well as the PVC interneuron, suggesting a pleiotropic effect on the nervous system. By comparing the genomes, transcriptomes and metabolomes of the most and least protective *E. coli* strains, we found that GABA is a key bacterial neuroprotective metabolite. Systemic *C. elegans* GABA receptors GAB-1 and LGC-37, and GABA transporter UNC-47 are required for wild type neuroprotection conferred by HT115 bacteria. Importantly, HT115 neuroprotective effect on neurons requires the function of the DAF-16 transcription factor.

### Gut microbes regulate neurodegeneration in *C. elegans*

Recent work has highlighted the importance of the gut microbiota in shaping human health and well-being. Not only do intestinal microbes regulate many aspects of host physiology and development, but they have also been linked to mood disorders and neurodegenerative diseases [7, 31]. For example, Parkinson Disease (PD) patients present constipation as non-motor symptom in early stage of the disease, which correlates with dysbiosis of the gut microbiome. Although it is unclear whether alterations in the gut microbiome are causes or consequences of these illnesses in the nervous system, emerging evidence using fecal transplantation in animal models demonstrate the ability of health and disease gut microbiome to ameliorate symptoms and confer disease, respectively [32, 33].

Intestinal microbiota regulates central nervous systems (CNS)-related traits through the microbiota-gut-brain axis. This consists in a bidirectional communication between the microbiota in the intestinal tract and the brain through the production of neuroactive molecules. Few recent studies correlate bacterial taxa from mammal’s microbiota with metabolite production and its impact on brain function and pathology (reviewed in ref. 34).

The bacterivore nematode *C. elegans* offers an advantaged platform for dissecting specific neuroprotective metabolites, because bacteria can be given monoaxenically to animals that express a genetically encoded degenerative trigger (*mec-4d*). Dietary bacteria are usually broken up in the grinder located in the worm’s pharynx right after ingestion [35]. This releases bacteria contents into the gut of the worm, a process that may also occur by explosive lysis [36, 37] Nonetheless, a number of dietary bacteria survive this initial interaction to live and colonize the worm’s digestive tract [38]. We show that UV-killed *E. coli* HT115 is equally neuroprotective as live bacteria, ruling out that bacteria need to colonize the intestine and actively trigger a host immunoprotective response. Moreover, HT115 fed only for a few initial hours to newly hatched animals was sufficient to provide protection to adult neurons. This suggests that HT115 metabolites turn on a signaling cascade that outlasts the presence of the protective bacterial metabolite itself.

### Bacterial GABA is neuroprotective

The glutamate decarboxylase system is a key mechanism used by intestinal bacteria to cope with acidic stress. This enzyme that produces GABA from glutamic acid is uniquely and highly expressed in HT115. GABA can be metabolized to succinic semi-aldehyde (SSA), and succinate; or translocated to the periplasm by the glutamate/GABA antiporter (GADT, [39]). GADT is dramatically overexpressed in protective bacteria likely tilting the balance towards the accumulation of GABA in the periplasm. Unbiased metabolomic analysis identified GABA as one of the main metabolites differentially expressed in *E. coli* HT115 compared to OP50. Finally, GABA supplementation to *E. coli* OP50 is sufficient to provide neuroprotection.

Some human enteric microbiota member have been shown to require GABA to grow, and thus the production of GABA by enteric members of the *Bacteroides, Parabacteroides* and *Escherichia* genera has been suggested to delineate the composition of the human microbiota [10]. Moreover, in the same work it was shown that the relative abundance of GABA-producing members in intestinal microbiota negatively correlates with depression-associated brain signatures in patients., Bacterial GABA was proposed as one of the main effectors of microbiota on the CNS [8-10]. How bacterially produced GABA in the gastrointestinal tract may affect the brain or other distal CNS traits is intriguing. In animals, GABA receptors are expressed in epithelial cells and a limited capacity of GABA to cross the blood-brain barrier has been reported (reviewed by [40]). However, is not clear if GABA and other microbiota produced neuroactive molecules may exert their effect by localized or systemic activation of signaling pathways. In *C. elegans* the number and type of GABA-containing cells and cells expressing GABA uptake proteins and receptors are higher than previously thought and includes non-neuronal cells [41]. Thus, although no intestinal GABAergic cells were reported, GABA could be pleiotropically sensed by a number of cells. In an infection model, *Staphylococcus aureus* molecules can trigger neuroendocrine reactions in *C. elegans* [42]. In our work, by using systemic and TRN specific reverse genetics, we show that systemically delivered dsRNA of *unc-47, lgc-37* and *gab-1* decreases neuroprotection in HT115 bacteria. This shows that GABA can be processed in non-neuronal cells to promote neuroprotection.

### DAF-16/ FOXO loss abolishes *E. coli* HT115 protection

A phosphorylation cascade downstream of the insulin receptor DAF-2/IGF1R controls DAF-16 transcription factor activation. When DAF-2/IGF1R is activated DAF-16/FOXO is prevented from entering the nuclei [26]. Starvation, pathogen exposure and other interventions promote its nuclear translocation and activation of transcriptional targets [43]. DAF-16 can also be directly activated upon fungal infection in a DAF-2 independent fashion [44]. In our experiments DAF-2 inactivation did not increase HT115 protection, suggesting they act in the same pathway. Neuroprotective and neuroregenerative effects of DAF-2 downregulation involve the function of DAF-16 [16, 22, 45]. Comparison of DAF-16 nuclear translocation between diets showed that at 24 hours DAF-16 was increased in the nuclei of animals feeding on HT115 compared to OP50. Nuclear presence of DAF-16 returned to basal levels at 48 hours. This raises the possibility that DAF-16 activity during that window of time is sufficient for long-term protection, by means of the stability of its transcriptional targets. Strikingly, while a *daf-16* mutation did not cause TRN loss of integrity in wild type animals, it completely abolished the protection by HT115 diet on *mec-4d* mutants. Taken together, these results show that DAF-16 is critical under conditions of neuronal stress and necessary for the dietary protection mediated by *E. coli* HT115 to take place.

## Materials and Methods

### *C. elegans* maintenance and growth

Wild type (N2), and mutant strains TU2773 [*uIs31*(*P*_*mec-17*_*mec-17::gfp*); *mec-4d(e1611)*], CF1139 [*daf-16(mu86)*; *muIs61*(*P*_*daf-16*_*daf-16::gfp*); *rol-6(su1006)*], WCH34 [*daf-2ts(e1368)*; *mec-4d(e1611); uIs31 (P*_*mec-17*_*mec-17::gfp*)], WCH39 [*daf-16*(*m27*); *mec-4d (e1611)*; *uIs31*(*P*_*mec-*_ *17mec-17::gfp*)], WCH40 [*daf-16*(*m27*); *uIs31*(*P*_*mec-17*_*mec-17::gfp*)]; TU38 [*deg-1(u38)*]; TU3755 [*uIs58(P*_*mec-4*_*mec-4::gfp)*]; WCH6 [*uIs71*(*P*_*mec-18*_*sid-1;Pmyo-2mcherry*), *uIs31*(*Pmec-17mec-17::gfp*), *sid-1*(*pk3321*), *mec-4d*(*e1611*)] were grown at 20°C as previously described [46] All nematode strains are maintained on *E. coli* OP50 strain prior to feeding with other bacteria. Unless otherwise noted, all plates were incubated at 20°C.

### Bacterial growth

Bacteria were grown overnight on Luria-Bertani (LB) plates at 37°C from glycerol stocks. The next morning a large amount of the bacterial lawn is inoculated in LB broth and grown for 6 hours on agitation at 450 g at 37°C. 100 ml of this bacterial culture is seeded onto 60 mm NGM plates and allowed to dry overnight before worms are placed on them. We used the following bacterial strains as worm food: *E. coli* OP50, *E. coli* HT115, *E. coli* K12, *E. coli* B, *Comamonas aquatica, Comamonas testosteroni, Bacillus megaterium*; *P. aeruginosa* PAO1, *Pseudochrobactrum sp, Stenotrophomonas sp, Bacillus pumilus*, and *E. coli* HT115 Δ*gad*.

### UV killing of bacteria

#### Killing on plates

300 μl of bacterial cultures grown at an OD of 0.8 were inoculated on 60 mm NGM plates. Once the liquid dried (overnight) plates were placed upside down in the UV transilluminator (Cole-Palmer HP 312nm) for 5 minutes at high power. Synchronized animals were seeded onto the plates immediately after bacterial killing. An inoculum of the UV treated bacterial lawn was picked and streaked onto new LB plates to confirm bacteria were effectively killed.

#### Killing in liquid

5 mL of bacterial cultures grown at an OD of 0.8 were placed in an empty 60 mm plate and exposed to UV light during 10 minutes with slow agitation every 5 minutes. 300 μl of this liquid is used to seed 60 mm plates.

### Criteria for neuronal integrity

AVM neuron: Morphological evaluation of AVM neuron was modified from Calixto et al., 2012 [16]. Neurons with full-length axons as well as those with anterior processes that passed the point of bifurcation to the nerve ring were classified as AxW (see Figure 1A). Axons with a process connected to the nerve ring were classified as AxL, and those that did not reach the bifurcation to the nerve ring as AxT. Lack of axon, only soma and soma with only the ventral projection was classified as AxØ.

ALM and PLM neurons: Neurons with full-length axons were classified as AxW. AxT were ALM neurons with axons that did not reach the bifurcation to the nerve ring and PLMs were axons that did not reach mid body. Neurons without axons or somas were classified as AxØ.

### Microscopy

For morphological evaluation, worms are mounted on 2% agarose pads, paralyzed with 1 mM levamisole, and visualized under a Nikon Eclipse Ti-5 fluorescence microscope with 40x or 60x magnification under Nomarski optics or fluorescence. For high-resolution images we used a Leica TCS SP5X microscope. DAF-16 nuclear expression and MEC-4 localization in CF1139 and TU3755 animals respectively were quantified using ImageJ (1.46v). For accuracy in the categorization and to avoid damage due to long exposure to levamisole animals were scored within 20 minutes after placing them on the agarose pads.

### Feeding RNAi

Clones of interest were taken from the Ahringer RNAi bacterial library [47, 48]. We performed a P0 screen as described in Calixto et al. [16] in *mec-4d* animals and TRN-autonomous RNAi strain carrying the *mec-4d*(*e1611*) mutation (WCH6, [16]. Bacterial clones were taken from glycerol stocks and grown overnight on LB plates containing tetracycline (12.5 µg/mL) and ampicillin (50 µg/mL). Next morning, a chunk of bacterial lawn was grown on liquid LB containing ampicillin (50 µg/mL) overday for 8 hours. 400 µL of bacterial growth was plated on NGM plates containing 1 mM IPTG and carbenicillin (25 µg/mL) and allowed to dry until next day 30-50 newly hatched *mec-4d* or WCH6 worms were placed on NGM plates. ALM integrity was scored in young adults 72 hours later.

### Synchronization of animals

Plates with large amounts of laid eggs were washed with M9 to eliminate all larvae and adults. Within the next two hours, newly hatched L1 animals were collected with a mouth pipette and transferred to the desired experimental plates.

### Time course of neuronal degeneration

Synchronized L1 larvae were placed in plates at 20°C with the desired bacterial food using a mouth pipette. We scored the integrity of the AVM neuron axon a) during development: at 12, 24, 48 and 72 hours post hatching, b) every 24 hours for longer periods (from 0 or 12 hours until 168 hours post hatching), and c) at adulthood (72 hours) after each treatment. The same experiments at 25°C also included ALM and PLM neurons. For experiments at 25°C without temperature shifts, parentals of animals examined were kept at the same temperature starting as L4s. Animals were scored until 168 hours at intervals of 24 hours. For each evaluation we used at least three biological replicas with triplicates of 30 worms each.

### Food changes

Synchronized L1 animals were placed in NGM plates seeded with UV killed *E. coli* HT115 or OP50 for 6 or 12 hours and later changed to *E. coli* OP50 or HT115 until 72 hours post hatching. To transfer 6- and 12-hour old animals to the new bacteria, larvae were washed off from the plates with sterile water supplemented with carbenicillin (25 µg/mL). Each replica was collected using a pipette in an Eppendorf tube. Animals were subsequently centrifuged for two minutes at 450 g. The pellet was washed with sterile water supplemented with carbenicillin followed by centrifugation. After two washes the pellet was collected and transferred to new plates. Axonal morphology was evaluated at 12, 24, 48 and 72 hours. For controls in both experiments, we grew synchronized animals in the same way on *E. coli* OP50 and HT115 for their entire development until 72 hours.

### DAF-16 expression on different foods

30 L4 CF1139 [*daf-16(mu86)*; *muIs61*(*P*_*daf-16*_*daf-16::gfp*); *rol-6(su1006)*] animals were placed in plates seeded with *E. coli* OP50 or HT115 and allowed to lay eggs for 24 hours. 30 synchronized 0-2 hours post-hatching L1 larvae were transferred with a mouth pipette to new plates with *E. coli* HT115 or OP50. Morphological evaluation was performed at 12, 24 and 48 hours post hatching.

### MEC-4 puncta quantification at different temperatures

30 L4 TU3755 [*uIs58 (P*_*mec-4*_*mec-4::gfp)]* worms were fed *E. coli* OP50 or HT115 and allowed to lay embryos for 24 hours. 30-50 L1 were synchronized as described above and placed on the corresponding diet at 15°C, 20°C or 25°C.

### Criteria for DAF-16 nuclear expression

We counted the number of GFP positive nuclei between the terminal pharyngeal bulb and 30 μm after the vulva (or developing vulva). Because we performed a time course evaluation, animals had different body sizes at each time point. The comparisons were made between animals of the same life stage feeding on the two bacteria. For 12-hour-old animals’ pictures were taken in a 60x objective and for 24 and 48-hour old animals, in a 40x objective. Experiments were done in triplicates for each time point.

### MEC-4 puncta quantification

One PLM per L4 animal were photographed under a 40X objective. Number of puncta was counted in 100μm of axon starting from the neuronal soma. Images were visualized in ImageJ (1.46v) and puncta counted manually.

### Touch response

To evaluate the functionality of the AVM mechanoreceptor neuron and PVC interneuron, the ability of animals to respond to gentle touch was tested. Animals were touched at 20°C.

First, animals were synchronized in L1 larvae and placed 30 per NGM plate seeded with different bacteria (mentioned above). Then, in the case of AVM neuron, animals were touched with an eyebrow one time in the head, gently stroking where the pharyngeal bulb lies at 72 hours, while PVC interneurons were gently touched with an eyebrow 10 times in a head to tail fashion every 24 hours after hatching.

### Temperature shifts

To evaluate the effects of DAF-2 downregulation on dauer animals, we used the strains WCH34 (*daf-2ts(e1368); uIs31 (P*_*mec-17*_*mec-17::gfp); mec-4d(e1611)* and TU2773 [*uIs31(P*_*mec-17*_*mec-17::gfp);mec-4d(e1611)*] as a control. L4 animals from both strains, were placed on plates seeded with *E. coli* HT115 and OP50 at 25 and 20°C. Then, animals were synchronized taken 30-50 L1 larvae and placed in three plates for each point of evaluation. Morphology of Touch receptor neurons (AVM, PLMs, ALMs and PVM) was scored from 12 to 168 hours with intervals of 24 hours.

### Longitudinal analysis of neuronal degeneration

30 synchronized L1s were placed on individual plates at hatching (0-2 hours) seeded with *E. coli* HT115. Animals were examined every 24 hours for 3 days, starting at 24 hours post hatching by placing them on 2 µl of 0,1 µm polystyrene beads for AVM observation and photography. We used polystyrene beads in order to maintain the shape of the animal, allowing its rescue from the agarose pad for posterior visualizations. Each animal was gently returned in M9 to the plate with a mouth pipette.

### Bacterial growth with controlled optic density (OD)

*E. coli* HT115 bacteria were inoculated in LB starting from a -80°C glycerol stock and allowed to grow for an hour. 9 different falcon tubes were used to grow bacteria. After the first hour, 1 mL from each tube was taken to measure the OD using a spectrophotometer (Ultraspec 2100). When the cultures reached the desired OD, growth was stopped. This procedure was repeated for each measurement. To obtain the lowest values (0.4/0.6/0.8), cultures were evaluated every hour, and for the rest of the values (0.8 to 2.0), every 15 or 30 minutes. 200 µL of each culture were inoculated onto six NGM plates. Seeded plates were dried on a laminar flow hood for at least 1 hour. Finally, bacteria were killed on a UV transilluminator (Cole-Palmer High performance) by placing the plate open upside down for 5 minutes, using the highest wavelength (365 nm). On three of those plates, 30 synchronized L1 worms were placed. The other three plates were used to test whether bacteria were killed during the protocol. To this end, parts of the lawn of UV-killed bacteria were streaked into another LB plate to observe if within the next three days, bacteria grew on the agar.

### Supplementation with bacterial supernatant

5 mL of overnight *E. coli* OP50 and HT115 bacterial cultures were centrifuged for 15 min at 3500g. The supernatant was sieved twice on 0,2 µm filters using a sterile syringe, to separate bacteria in suspension. *E. coli* OP50 supernatant was used to resuspend *E. coli* HT115 pellet, and the same was done with *E. coli* HT115 supernatant on OP50 pellet. Each bacterial suspension was inoculated on NGM plates as described above.

### Supplementation with GABA or glutamic acid

After liquid UV-killing, the desired bacteria were mixed with 2 mM γ-aminobutyric acid, or 2 mM glutamate (Sigma). 400 μl of bacterial cultures was inoculated in 60 mm NGM plates to cover the entire surface. Next day 0-2 hour synchronized L1 worms were placed in the NGM plates for 24 hours. Animals were moved to freshly prepared plates every 24 hours until 72 hours.

### Mix of bacteria and UV killing

Bacterial cultures were combined in different proportions with the other bacteria or LB for controls. Liquid LB media growth until OD 0,6 of *E. coli* OP50 or HT115 were diluted 1:125 in LB and incubated at 37°C with shaking O/N. Bacterial cultures were mixed in the indicated proportions and used to prepare plates as described before (0.1, 1, 10, and 50 % of *E. coli* HT115 in OP50). Plates with mixed bacteria were UV-irradiated before adding the worms.

### Generation of bacterial *gad* mutant

Two orthologs of the glutamate decarboxylase gene are present in *E. coli* HT115, denominated *gadA* and *gadB. E. coli* HT115 mutants were constructed using homologous recombination with PCR products as previously reported [49]. Wild type *E. coli* HT115 [50] was transformed with plasmid pKD46 [49]. Next, electrocompetent bacteria were prepared at 30°C in SOB medium with ampicillin (100 µg/ml) and arabinose and electroporated with a PCR product obtained using the set of primers gadBH1P1 and gadBH2P2 and pKD3 as template. Recombinant candidates were selected in LB plates plus chloramphenicol at 37°C. Colonies were tested for loss of ampicillin resistance. Amp^s^ colonies were checked for substitution of the *gadB* gene by the chloramphenicol acetyl transferase cassette by PCR using primers gadB-A and gadB-B, which flank the insertion site. This rendered the *E. coli* HT115 Δ*gadB*::*cat* derivative.

This strain was electroporated with pKD46. Competent cells of this transformed strain were prepared with ampicillin and arabinose and electroporated with a PCR product generated using primers NgadAH1P1 and NgadAH2P2 and pKD4 as template.

Recombinant candidates were selected in LB plates plus kanamycin (30 µg/ml) at 37°C. Colonies obtained were tested for loss of ampicillin resistance and checked for substitution of the *gadA* gene by the kanamycin resistance gene through PCR using primers NGadAFw and NgadARv, flanking the substitution site. This yielded an *E. coli* HT115 Δ*gadB*::*cat*/Δ*gadA*::*kan* double mutant strain (referred as *E. coli* HT115Δ*gad* mutant in the text).

### Generation of the pG*gadA* plasmid

*E. coli* HT115 *gadA* gene was amplified by PCR using primers NgadAFw and NgadARw with Taq polymerase and cloned in the pGemT Easy plasmid (Promega) according to manufacturer’s instruction, to generate pG*gadA. E. coli* OP50 and *E. coli* HT115 Δ*gadB*::*cat*/Δ*gadA*::*kan* were transformed chemically with the pG*gadA* plasmid. Competent bacteria were obtained using a calcium chloride protocol [51]. Transformed colonies for *E. coli* OP50+pG*gadA* and HT115Δ*gadB*::*cat*/Δ*gadA*::kan+pG*gadA* were confirmed by antibiotic resistance (ampicillin 100 µg/ml), plasmid purification, plasmid length and gene endonuclease restriction with XbaI (NEB).

### GAD enzymatic activity

Glutamate decarboxylase enzymatic activity was measured according to Rice et al. [28], with modifications [27]. Fresh bacterial colonies were grown in LB with the appropriate antibiotic until 108 CFU/mL (3-4 hours). Antibiotic used were 25 µg/ of streptomycin for *E. coli* OP50, 25 µg/mL of tetracycline for HT115, 20 µg/mL of kanamycin and chloramphenicol for HT115Δ*gad*, and 25 µg/mL of streptomycin and ampicillin and 25 µg/mL of IPTG for OP50+pG*gadA*.

Then, 10 mL of each culture were centrifuged at 500g for 10 minutes and the pellet was resuspended in 5 mL of phosphate buffer (KPO4 1M, pH 6.0). This step was repeated one more time and the pellet was resuspended in 2 mL of GAD reagent (1 g glutamic acid (Sigma), 3 mL triton X-100 (Winkler), 0.05g bromocresol blue (Winkler), 90 g sodium chloride (Winkler) for 1 liter of distilled water with final pH 3.4), preserved at 4°C for 2 months maximum. Once the pellet was resuspended the samples were measured for colorimetric differences (UltraSpect 2100) at 620nm. Liquid LB treated without bacteria is used as blank. Average colorimetric value for *E. coli* HT115 was considered 100% of possible GAD activity for each replicate.

### Growth of bacterial cultures for GABA quantification by GABase assay

Bacterial cultures were grown in LB media with 25 µg/mL of streptomycin for *E. coli* OP50, 25 µg/mL of tetracycline for HT115, 20 µg/mL of kanamycin and chloramphenicol for HT115Δ*gad*, until 10^7^ CFU/mL (3-4 hours).

### Bacterial GABA quantification by GABase assay

Measurement of GABA was performed following the GABase® Sigma protocol with modifications [29]. Two milliliters of each culture were centrifuged at 845 g for 10 minutes. Pellet and supernatant were separated, and supernatant received an additional centrifugation step at 3300 g. Then, pellets were washed with 2 mL of phosphate buffer (KPO_4_ 1M, pH 6.0). Samples were incubated in a boiling water bath for 15 minutes. Debris from cell lysis were centrifuged at 845 g for 10 minutes, and 0.16 mL of supernatant were placed in the GABase reaction for a final volume of 0.6 mL with the following final concentrations: 50 mM potassium pyrophosphate, 18 mM of dithiothreitol, 1.25 mM of β-NADP, 1 mM of potassium phosphate, 0.33 % (v/v) of glycerol and 0.02 units of GABase. A linear correlation among absorbance and GABA concentration was fitted to the equation y=0.102 x + 0.0254, with a correlation coefficient r2= 0.9957, and GABA levels were measured in the range of 0.01 to of 0.7 mM during a 2 hours reaction.

This reaction consisted in two enzymes that convert a molecule of GABA to succinate by GABA transaminase (GAT) and succinic semialdehyde dehydrogenase (SADH), producing detectable NADPH at 340 nm. Additionally, by inhibition of GAT with aminoethyl hydrogen sulfate, substrates concentrations for each reaction are distinguishable according to: OD [NADPH] total = OD [NADPH] GAT + OD [NADPH] SADH.

### Growth of bacteria for NMR Spectroscopy

Bacterial strains were grown in solid LB media with 25 µg/mL of streptomycin for *E. coli* OP50, 25 µg/mL of tetracycline for *E. coli* HT115, 20 µg/mL of kanamycin and chloramphenicol for HT115Δ*gad*, at 37°C for 10 hours. Eight pre-inoculums of 2 mL of liquid LB were set for each bacterial strain, grown overnight and continue in 35 mL of LB and monitoring by OD until required (OD of 1).

### Metabolite extraction for NMR

Bacterial cultures were centrifuged at 4000g for 5 minutes to separate bacteria from the media. Bacterial pellet was resuspended in phosphate buffer saline (PBS: NaCl (137 mM), KCl (2.7 mM), Na2HPO4 (10mM) and KH2PO4 (1.8 mM)) and washed twice after centrifugation with PBS at 4000g for 5 minutes. After the last wash each pellet was resuspended in 1 mL of cold extraction buffer and pipetted into Eppendorf tubes. The extract solution was prepared by mixing equal volumes of acetonitrile and K2HPO4/NaH2PO4, 100 mM, pH 7.4.

Bacterial membrane permeabilization was performed in two steps. First, Eppendorf tubes were submerged in a liquid nitrogen bath for 2 minutes, defrosted at 4°C and vortexed for 30 seconds. This procedure was repeated three times. Secondly, tubes from the first step were sonicated in an ultrasonic water bath (Bioruptor UCD-200, Diagenode) for 15 cycles of 30 seconds on and 30 seconds off at full power. Finally, to obtain the metabolites samples were centrifuged for 10 minutes at 8000 g and the supernatant was recovered. This step was repeated to optimize metabolite recovery. Samples were dried in a vacuum dryer (Savant) for 60min at 50°C and 300 g.

#### Metabolic Profiling

1H-NMR spectroscopy and multivariate data analysis were performed at PLABEM (Plataforma Argentina de Biología Estructural y Metabolómica).

### Sample Preparation for 1H NMR Spectroscopy

Samples were randomized and reconstituted in 600 µL of 100 mM Na^+^/K^+^ buffer pH7.4 containing 0.005% TSP (sodium 3-trimethylsilyl- (2,2,3,3-2H4)-1-propionate) and 10% D_2_O. In order to remove any precipitate, samples were centrifugated for 10 min at 14300 g at 4°C. 500 µL of the centrifuged solution were transferred into 5 mm NMR tube (Wilmad LabGlass).

### 1H NMR Spectroscopic Analysis of Bacterial Extracts

NMR spectra were obtained at 300 K using a Bruker Avance III 700 MHz NMR spectrometer (Bruker Biospin, Rheinstetten, Germany) equipped with a 5 mm TXI probe. One-dimensional 1H NMR spectra of bacterial extracts were acquired using a standard 1-D noesy pulse sequence (noesygppr1d) with water presaturation [52, 53], The mixing time was set to 10 ms, the data acquisition period to 2.228 s and the relaxation delay to 4 s. 1 H-NMR spectra were acquired using 4 dummy scans and 32 scans, with 64K time domain points and a spectral window of 20 ppm. FIDs were multiplied by an exponential weighting function corresponding to a line broadening of 0.3 Hz. Two-dimensional NMR spectra

### Quality controls

Quality controls (QC) were prepared as suggested by Dona et al. [52]. 1H NMR spectra of QC samples were acquired every 8 study samples.

### 1H NMR Spectral Processing

Spectroscopic data was processed in Matlab (Version R2015b, The MathWorks Inc., United States). Spectra were referenced to TSP at 0.0 ppm, baseline correction and phasing of the spectra was achieved using Imperial College written functions (provided by T. Ebbels and H. Keun, Imperial College London). Each spectrum was reduced to a series of integrated regions of equal width (0.04 ppm, standard bucket width). Non-informative spectral regions containing no metabolite signals, TSP signal and the interval containing the water signal (between 4.9 and 4.6 ppm) were excluded. Each spectrum was then normalized by probabilistic quotient method [54]. Spectra alignment was made using the alignment algorithm recursive segment-wise peak alignment (RSPA, [55]) in user-defined windows.

### Statistical Analysis of NMR Spectroscopic Data

The pre-processed 1H NMR spectral data was imported to SIMCA (version 14.1, Umetrics AB, Umeå, Sweden) for multivariate data analysis. Principal Component Analysis (PCA) was performed on the Pareto-scaled NMR dataset. Orthogonal partial least squares discriminant analysis (OPLS-DA) was made to maximize the separation between bacterial groups as function of neuroprotection. To ensure valid and reliable OPLS-DA models and to avoid overfitting, 200 permutations were carried out. Discriminant features between classes in OPLS-DA models were defined using a combination of Loading plot (S-line plot) and VIP plot. Variables met highest p(corr)1 in S-Line plot and VIP values□>1□were selected and validated by spectral raw data examination.

### NMR Resonances Assignment

Two-dimensional NMR spectra 1H J-resolved pulse sequences were acquired for resonance assignment purposes. Discriminant features were assigned searching in ¨The *E. coli* metabolome database¨ (ECMDB) [56, 57] The unequivocal identification of GABA and glutamate were made through spike-in experiments over an *E. coli* HT115 and HT115Δ*gad* extract, respectively.

### Growth of bacteria expressing *gad* plasmid

Genetically complemented bacteria with pG*gadA* were grown overday until OD of 0.6. GAD expression was induced by adding 0.15 mM IPTG to OD 0.6 liquid cultures. After 1-hour samples of each condition were taken to assess GAD activity and to seed on NGM plates with the appropriate antibiotic and supplemented with 0,1mM of IPTG.

### Genome sequencing of *E. coli* OP50 and HT115

*E. coli* OP50 and HT115 were grown from glycerol stock on LB plates overnight. Next morning portions of the lawn were cultured on agitation for 4 hours in liquid LB. 2 mL of liquid cultures were pelleted and DNA was purified using the UltraClean microbial DNA isolation Kit (MO BIO) according to manufacturer instructions.

Whole genome sequencing was done at Genome Mayor Sequencing Services. Paired-end reads (2×250bp) were generated using the Illumina MiSeq platform. Sequence data was trimmed using Trimmomatic version 0.27 [58]. Trimmed reads were assembled using SPAdes version 3.1.0 [59]. Genome annotation for both organisms was done using PROKKA version 1.9 [60]. Both genomes were deposited to NCBI under the accession numbers PRJNA526029 (*E. coli* OP50) and PRJNA526261 (*E. coli* HT115).

### Transcriptomic analysis

For total RNA isolation, *E. coli* OP50 and HT115 were grown from glycerol stock on LB plates overnight. Next morning portions of the lawn were cultured on agitation in liquid LB for 5 hours. Then, 2 mL of bacterial cultures were pelleted for RNA extraction with Max Bacterial Trizol kit® (Invitrogen) according to manufacturer protocol.

cDNA libraries for Illumina sequencing were generated by Centro de Genomica y Bioinformatica, Universidad Mayor, Chile. cDNA libraries were made with Illumina Truseq stranded mRNA kit, according to manufacturer protocol, in Illumina HiSeq plataform. Quality control of libraries was made with Bioanalyzer and quantification with qPCR Step One Plus-Applied Biosystem. Six sets of lllumina paired-end reads in FASTQ format corresponding to three replicates from *E. coli* strain HT115 and three replicates from strain OP50 were analyzed as follows.

#### Data pre-processing and quality control

Reads with an average quality lower than 30 over four bases, as well as reads shorter than 16bp were discarded with Trimmomatic version 0.35 [58]. Pre and post-trimming quality visualization was made with FastQC (https://www.bioinformatics.babraham.ac.uk/projects/fastqc/).

#### Identification of common and unique sequences of E. coli strains

Annotated proteins from assembled genomes were compared between strains by Reciprocal Best Hit analysis using Blast+ [61, 62]. We ran Blastp [63] of the protein sequences of OP50 strain against HT115 and selected the top hits (best match based on bit-scores and E-values). Then, the selected sequences from the *E. coli* HT115 strain were compared against OP50, and we extracted top hits. “Common sequences” are defined as those that were the best match of each other. All the other sequences were defined as “unique”.

#### Mapping and quantification of transcript abundance

Mapping and quantification from *E. coli* HT115 and OP50 strains were made using Salmon 0.12.0 [64] in quasi-mapping mode. For better comparability in transcript abundance between two slightly different strains we used the same reference. For this, we built a composite transcriptome by combining both common and unique sequences (defined in previous step) from both strains and removing redundant entries. *Differential expression analysis*. Transcript abundance was compared between strains. For this, Salmon quantification output was imported for subsequent analysis in R (version 3.5) with tximport package [65] then, differential expression analysis was performed using DeSeq2 version 3.8 [66] using default parameters. Cutoff for differentially expressed genes (DEG) was set at adjusted p value (padj) < 0.05, and are reported in Supplementary File 2.

#### Criteria for gene expression level

We categorized gene expression according to [67] as: low if expression is between 0.5 to 10 FPKM or 0.5 to 10 TPM; medium if expression is between 11 to 1000 FPKM or 11 to 1000 TPM; and high if expression was more than 1000 FPKM or 1000 TPM.

Detailed pipeline for transcriptomic analysis decribed above is available at the following link: https://mlegue@bitbucket.org/mlegue/ht115_op50.git

### Biological and technical replicates

Each experiment was performed in three technical triplicates and at least three biological replicates. We define biological replicates as experiments made in different days, containing triplicates of each condition, and a technical replicate as a triplicate of the same condition on the same day. The average of the three reads of each triplicate is considered as one count. Each experiment has three technical replicates that were in turn averaged to constitute one of the points of each figure. Data is collected and processed as a single technical replicate (the average of three counts of the same plate), and its mean is used as a single biological replicate. Each figure contains at least three experiments (biological replicates) performed as explained before. All the biological replicates are performed spaced from each other from 1 day to 1 week.

#### Sample size

Each experiment started with at least 30 synchronized worms on each technical triplicate, with exception of the longitudinal study of degeneration on *E. coli* HT115 that used 30 animals in total.

### Statistical evaluation

Statistical evaluation was performed using one or two way-ANOVA, with *post-hoc* tests, and a student t-test when indicated. Results of all tests are detailed in Supporting Information.

## Supporting information

Dataset all data

Dataset all statistics

differential expression analysis bacteria

Expression levels bacterial genes

List of metabolites

Genomics analysis unique genes

## Figure Legends

**Supplementary Table 1. Summary of number of shared or unique genes from *E. coli* OP50 and HT115 separated by expression level**.

**Supplementary Table 2. 1H NMR resonance assignments**

**A. *E. coli* HT115 and HT115** Δ***gad* mutant extracts.** S=singlet; d=doublet; dd= double doublet; t-triplet; m= multiplet. **B. Neuroprotective (HT115) and no neuroprotective (OP50 and HT115) bacterial extracts.** S=singlet; d=doublet; dd= double doublet; t-triplet; m= multiplet.

**Supplementary Figure 1.**
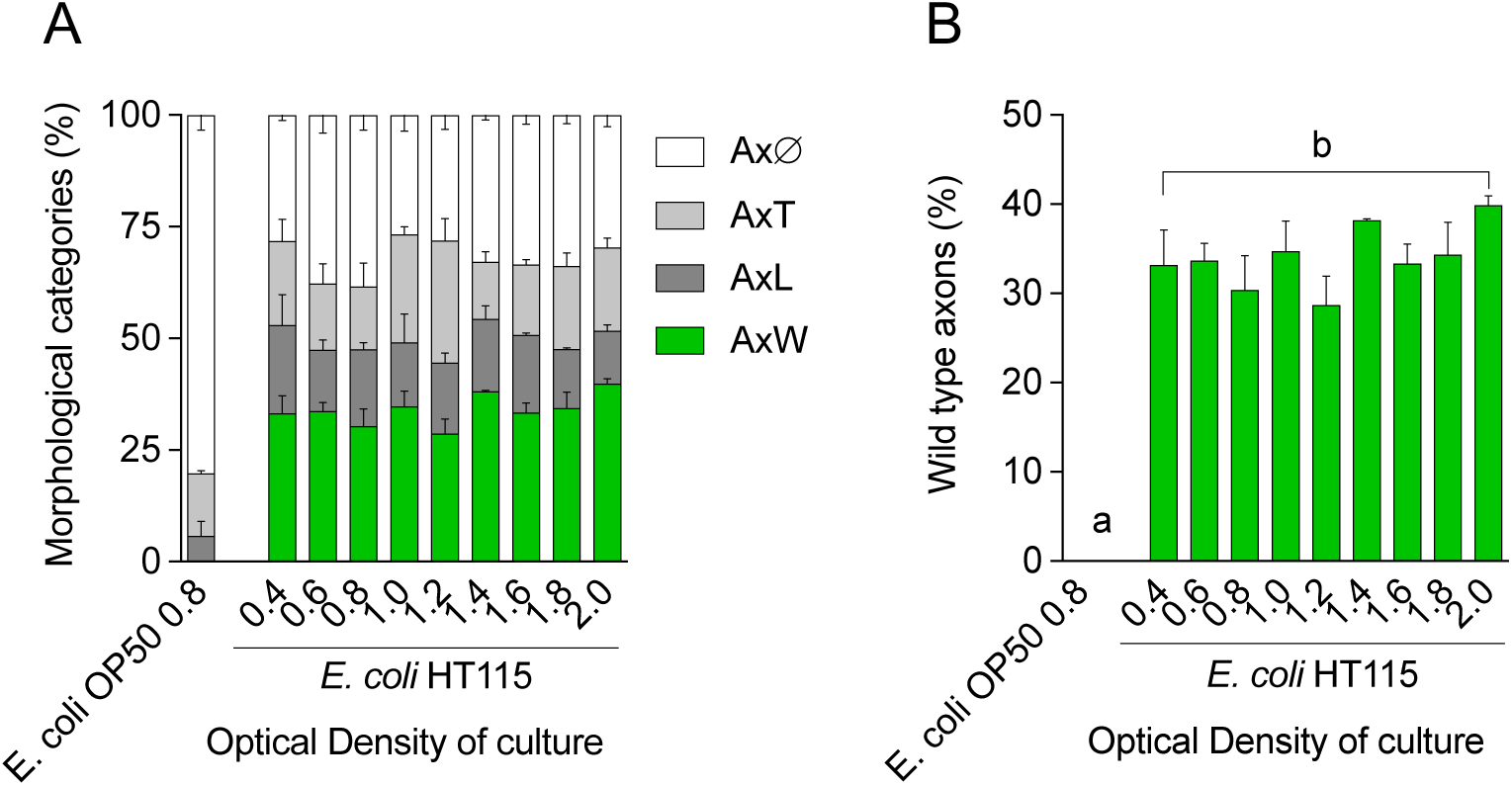
Axonal categories at different optical density. **A-B**. All axonal categories (**A**) and wild type axons (**B**) in worms feeding on *E. coli* HT115 bacteria grown to different optical density.

**Supplementary Figure 2.**
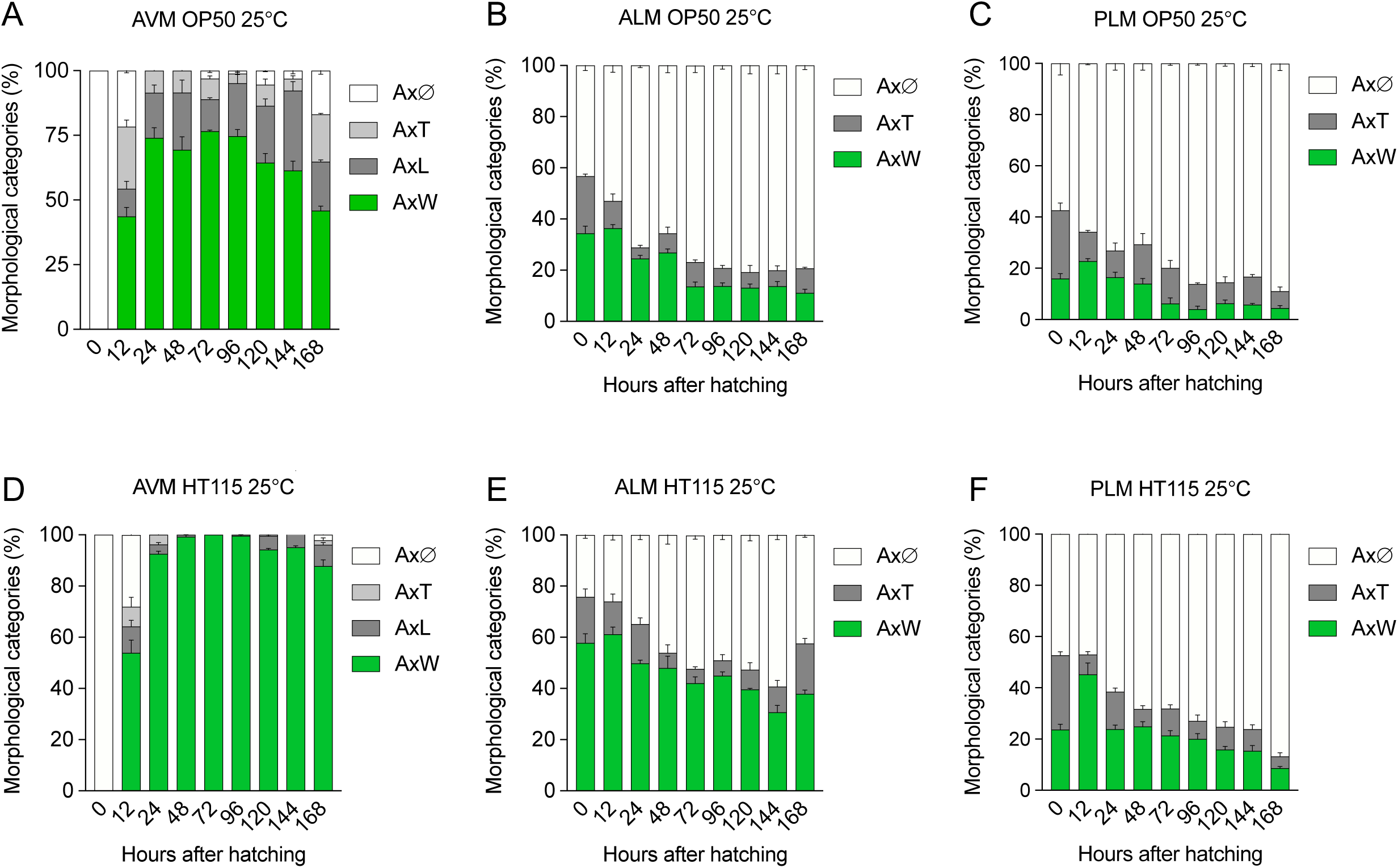
Axonal morphological categories of different classes of neurons in *E. coli* HT115 and OP50. All axonal categories of animals feeding on *E. coli* OP50 (**A, C** and **E**) and HT115 (**B, D** and **F**) at 25°C.

**Supplementary Figure 3.**
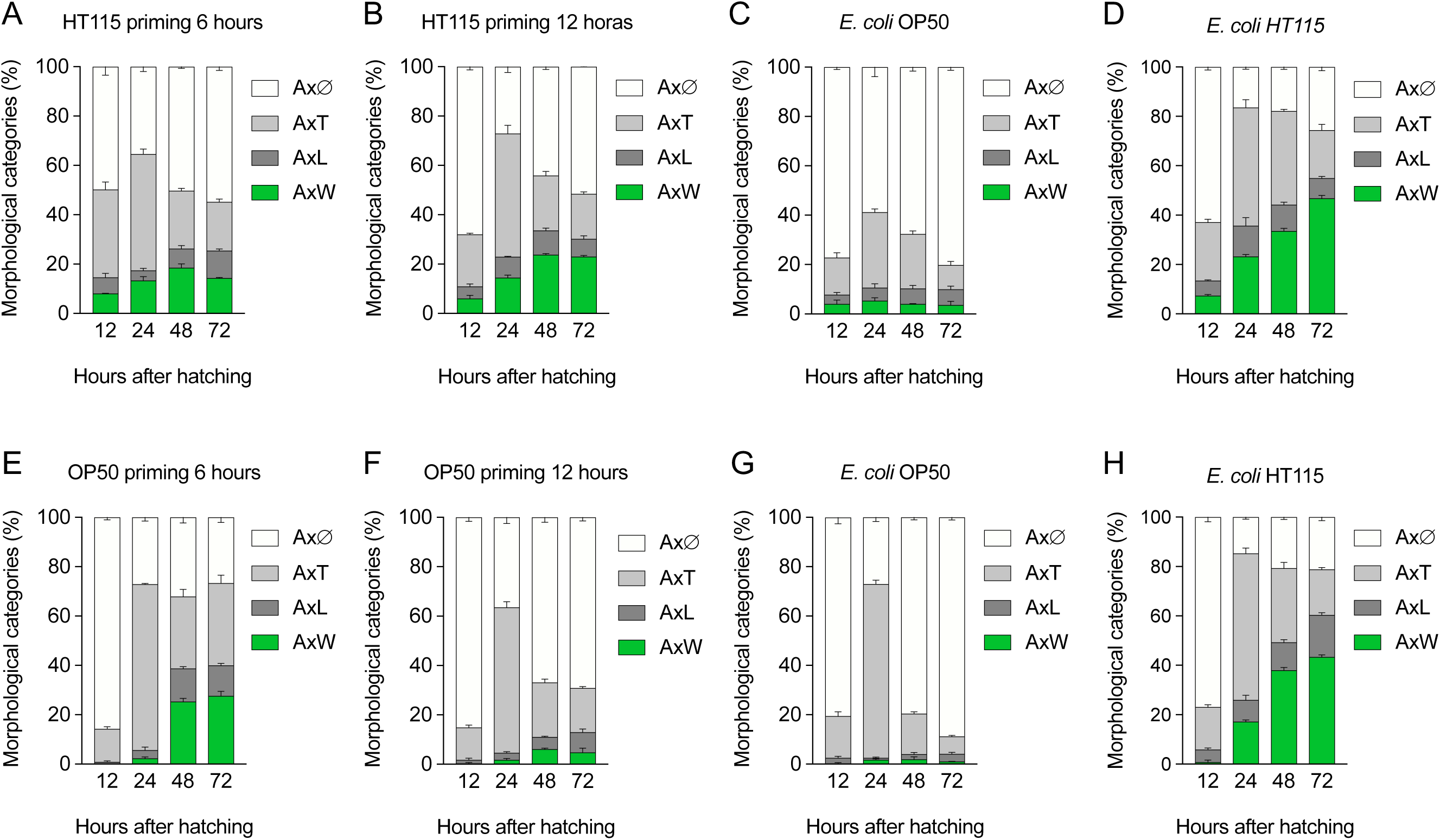
Complete axonal categories of priming experiments. **A-D**. All axonal categories of animals feeding *E. coli* HT115 for 6 (**A**) and 12 (**B**) hours with controls of *ad libitum E. coli* OP50 (**C**) and HT115 (**D**); or feeding *E. coli* OP50 for 6 (**E**) and 12 (**F**) hours with controls of *ad libitum E. coli* OP50 (**G**) and HT115 (**H**).

**Supplementary Figure 4.**
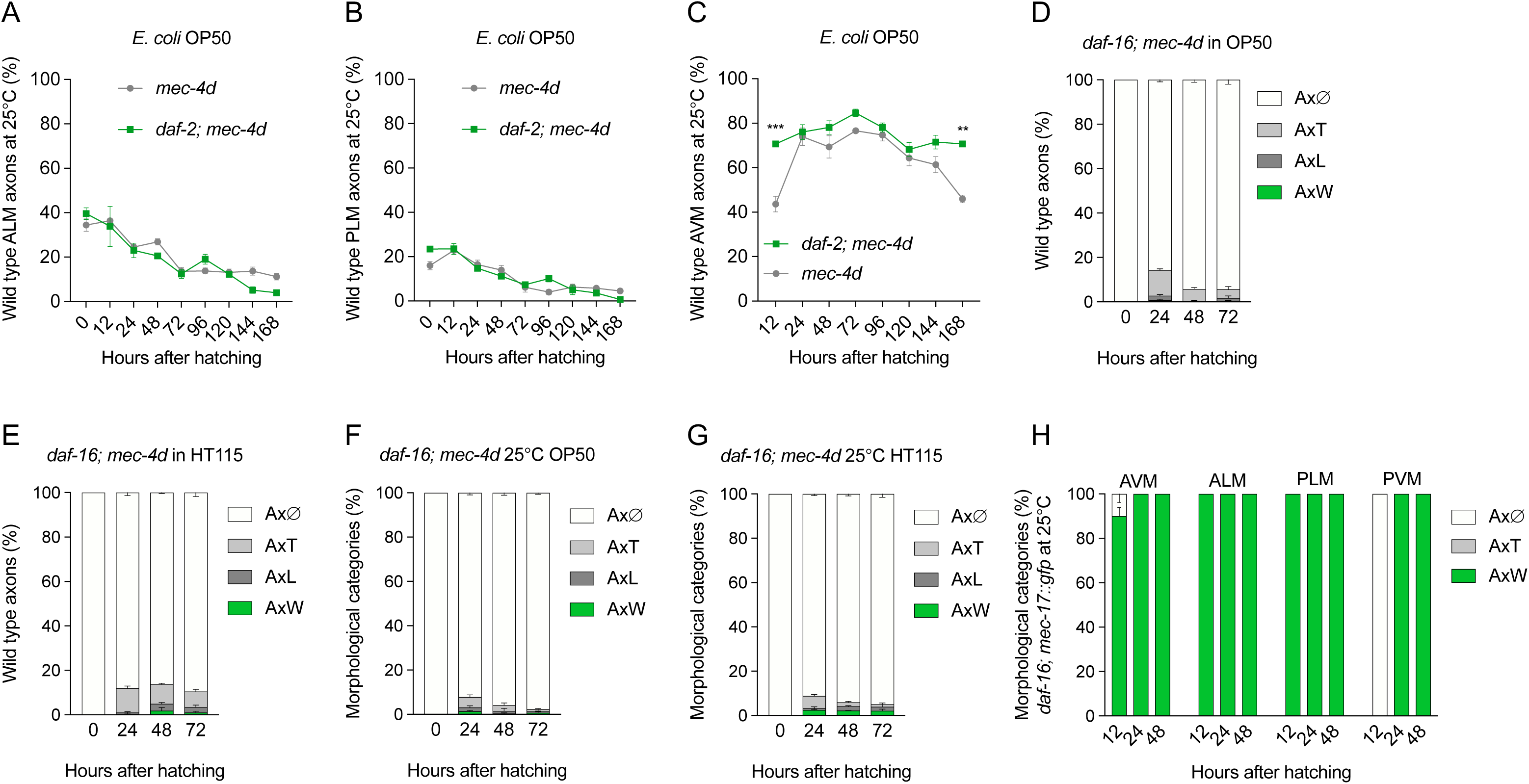
Effect of DAF-2 downregulation on neuronal degeneration in animals feeding *E. coli* OP50. **A-C** Neuronal integrity of AVM (**A**), ALM (**B**) and PLM (**C**) neurons of *daf-2(ts*); *mec-4d* animals feeding on OP50 food. D-E All axonal categories of *daf-16; mec-4d* animals feeding on *E. coli* OP50 (D) and HT115 (E).

**Supplementary Figure 5.**
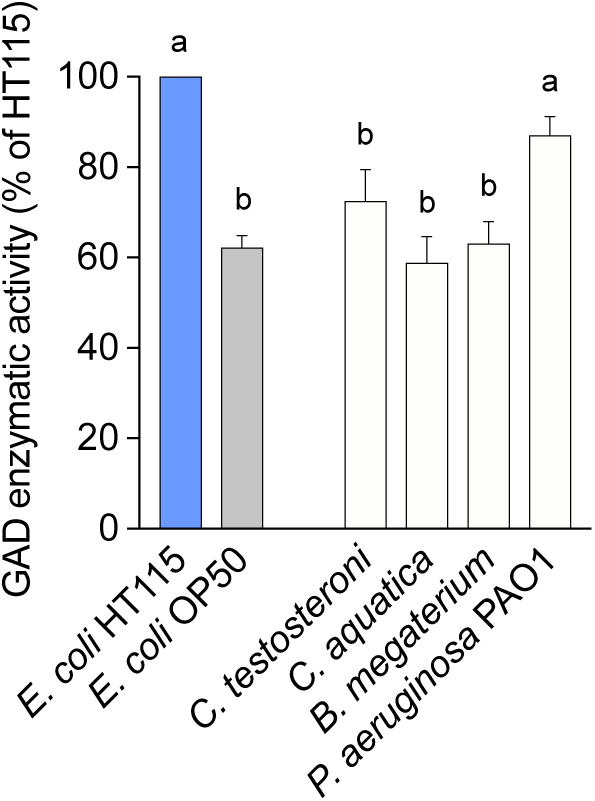
Measurements of GAD enzyme activity of normalized as a percentage of HT115. GAD activity in all strains tested for neuroprotection in Figure 1B.

**Supplementary Figure 6.**
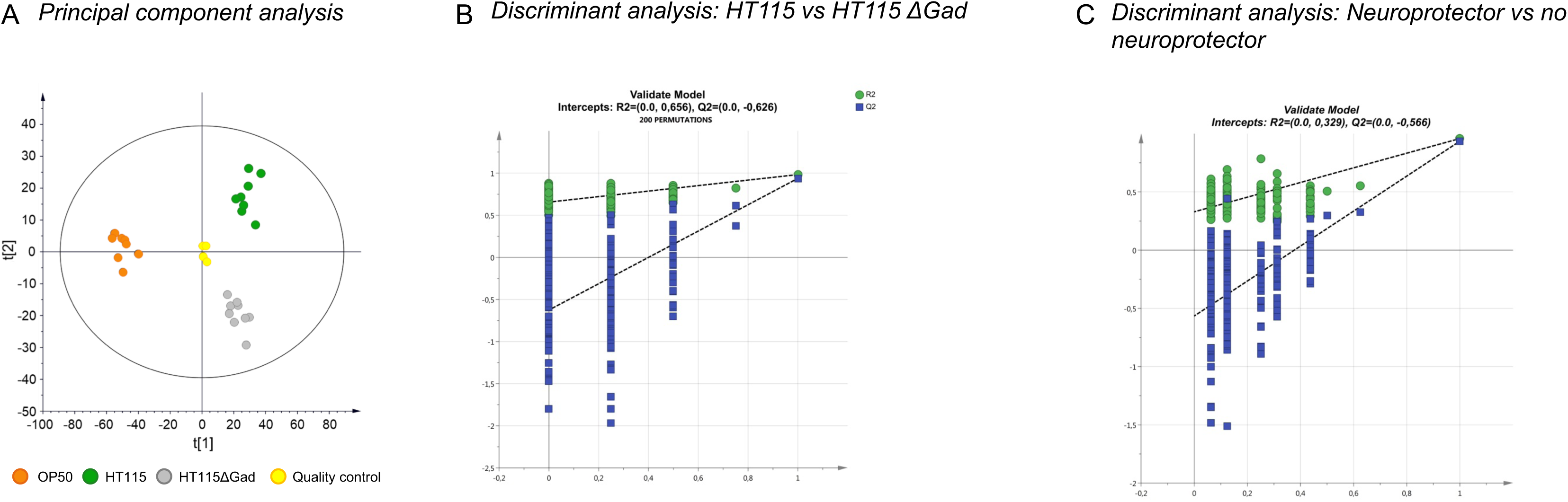
Multivariate analysis and models validation. **A.** Principal Component (PC) scores plot derived from 1H NMR spectra indicating metabolic differences between wild type *E. coli* strains; OP50 (orange) and HT115 (green) and HT115 Δ*gad* mutant (light gray). Quality controls are displayed in yellow. Model parameters are R2X = 0.787 Q2= 0.705. **B-C**. OPLS-DA Validation by 200 permutations. *E. coli* HT115 and HT115 Δ*gad* validate model intercepts: R2= (0.0; 0.656) and Q2= (0.0; -0.626) (**B**). Protective *E. coli* HT115 and non-protective strains *E. coli* OP50 and HT115 Δ*gad* validate model intercepts: R2= (0.0; 0.329) and Q2= (0.0; -0.566) (**C**).

**Supplementary Figure 7.**
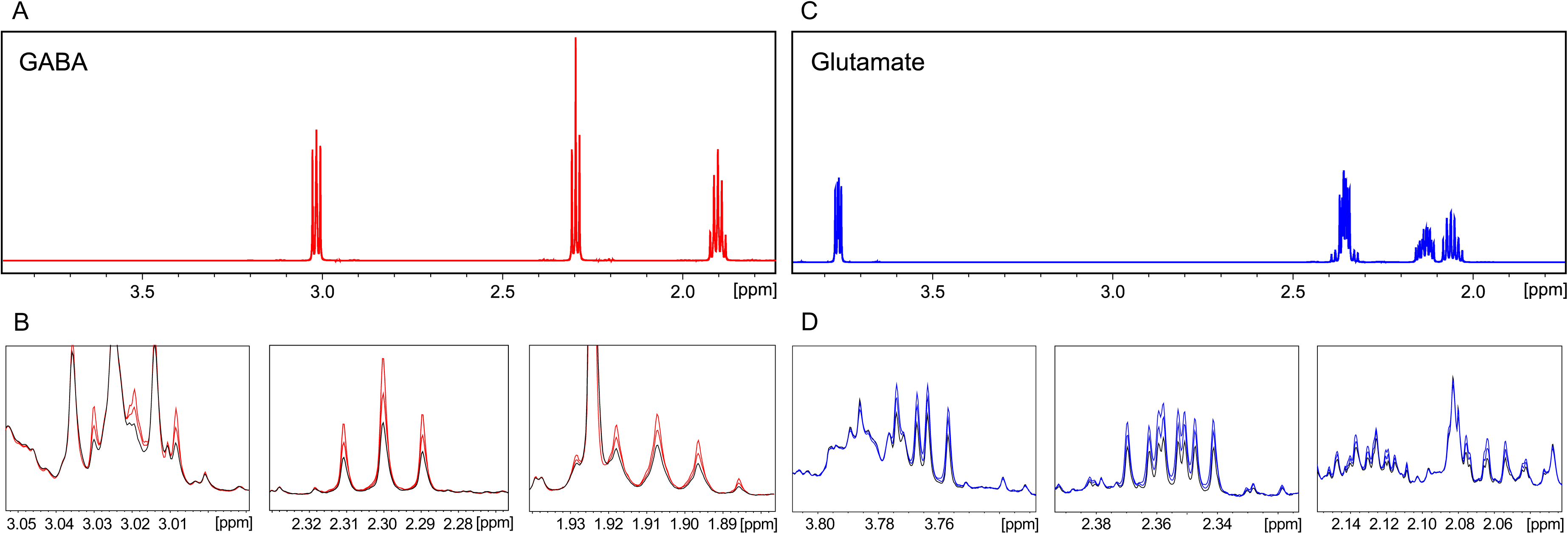
Resonances assignment confirmation by NMR. **A-C** 1H NMR spectra of GABA (**A**) and glutamate (**C**). **B-D** Spike-in of GABA and glutamate confirms identity of metabolites. *E. coli* HT115 extract (red) **(B)**. *E. coli* HT115 Δ*gad* extract (blue). Spike-in was made adding 5 µL of standard 10 mM twice.

**Supplementary Figure 8.**
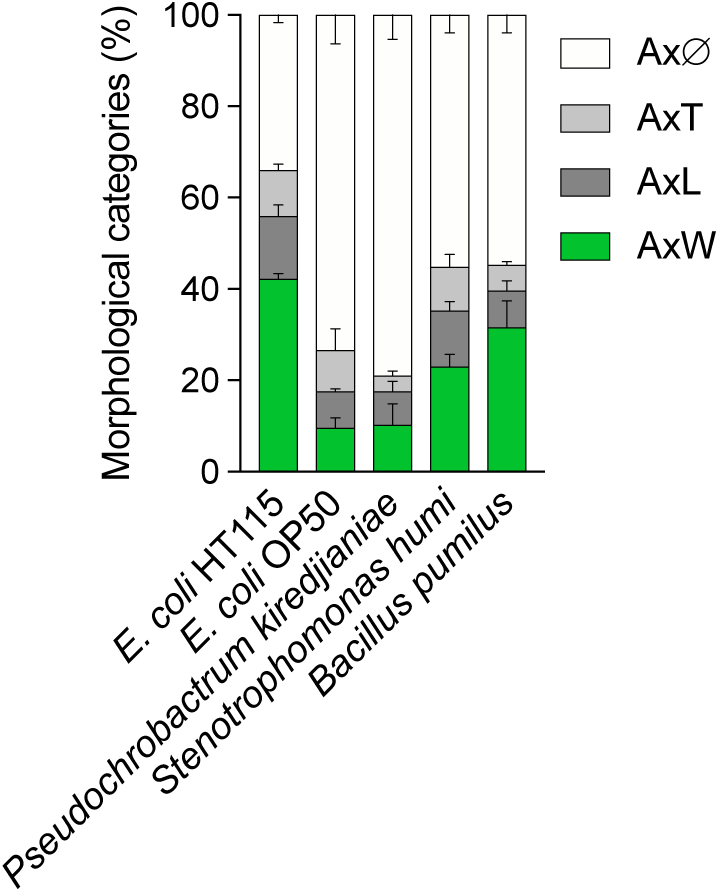
Complete axonal categories of animals feeding on soil bacteria. All axonal categories of animals feeding *Pseudochrobactrum kiredjianiae, Stenotrophomonas humi*, and *Bacillus pumilus* for 72 hours.

**Supplementary File 1.** Genomics data showing unique genes of *E. coli* OP50 and HT115.

**Supplementary File 2.** Differential expression analysis for all shared genes between *E. coli* OP50 and HT115

**S1 Dataset.** All data with replicas from the experiments shown in manuscript.

**S2 Dataset.** All statistical analysis of the experiments contained in the manuscript.

## Acknowledgements

We thank Irini Topalidou and Ines Carrera for critical reading of the manuscript and Diego de Mendoza for providing laboratory space and reagents. Some strains were provided by the CGC, which is funded by NIH Office of Research Infrastructure Programs (P40OD010440). The Centro Interdisciplinario de Neurociencia de Valparaiso is a Millennium Institute supported by the Millennium Scientific Initiative of the Chilean Ministry of Economy, Development, and Tourism (P029-022-F).

## Author contribution

Conceptualization A.C; Methodology P.B, A.U, V.A.G and A.C; Investigation A.U, V.A.G, A.F, M.C, P.B, M.L, and A.C.; Writing-Original Draft, A.C; Writing-Review and Editing, V.A.G, P.B and A.C; Funding Acquisition A.C.

